# Immunogenicity of an AAV-based, room-temperature stable, single dose COVID-19 vaccine in mouse and non-human primates

**DOI:** 10.1101/2021.01.05.422952

**Authors:** Nerea Zabaleta, Wenlong Dai, Urja Bhatt, Jessica A Chichester, Julio Sanmiguel, Reynette Estelien, Kristofer T Michalson, Cheikh Diop, Dawid Maciorowski, Wenbin Qi, Elissa Hudspeth, Allison Cucalon, Cecilia D Dyer, M. Betina Pampena, James J. Knox, Regina C LaRocque, Richelle C Charles, Dan Li, Maya Kim, Abigail Sheridan, Nadia Storm, Rebecca I Johnson, Jared Feldman, Blake M Hauser, Eric Zinn, Aisling Ryan, Dione T Kobayashi, Ruchi Chauhan, Marion McGlynn, Edward T Ryan, Aaron G Schmidt, Brian Price, Anna Honko, Anthony Griffiths, Sam Yaghmour, Robert Hodge, Michael R. Betts, Mason W Freeman, James M Wilson, Luk H Vandenberghe

**Affiliations:** Grousbeck Gene Therapy Center, Schepens Eye Research Institute, Mass Eye and Ear, Boston, Massachusetts, USA; Ocular Genomics Institute, Department of Ophthalmology, Harvard Medical School, Boston, Massachusetts, USA; The Broad Institute of Harvard and MIT, Cambridge, Massachusetts, USA; Harvard Stem Cell Institute, Harvard University, Cambridge, Massachusetts, USA; Gene Therapy Program, Perelman School of Medicine, University of Pennsylvania, Philadelphia, Pennsylvania, USA; Novartis Gene Therapies, San Diego, California, USA; Novartis Gene Therapies, North Carolina, USA; Department of Microbiology, Perelman School of Medicine, University of Pennsylvania, Philadelphia, Pennsylvania, USA; Department of Pathology, Perelman School of Medicine, University of Pennsylvania, Philadelphia, Pennsylvania, USA; Division of Infectious Diseases, Massachusetts General Hospital, Boston, MA; Department of Medicine, Harvard Medical School, Boston, MA; Department of Microbiology and National Emerging Infectious Diseases Laboratories, Boston University School of Medicine, Boston, MA 02118, United States; Ragon Institute of MGH, MIT, and Harvard, Cambridge, MA, 02139, USA; Translational Innovation Fund, Mass General Brigham Innovation, Cambridge, Massachusetts, USA; Department of Immunology and Infectious Diseases, Harvard T. H. Chan School of Public Health, Boston, MA; Department of Microbiology, Harvard Medical School, Boston, MA, USA; Albamunity, Boston, Massachusetts, USA; Novartis Gene Therapies, Libertyville, Illinois, USA; Center for Computational & Integrative Biology, Department of Medicine, and Translational Research Center, Massachusetts General Hospital, Harvard Medical School, Boston, Massachusetts, USA

**Keywords:** Adeno-associated virus, AAV, SARS-CoV-2, COVID-19, vaccine, immunization, single dose

## Abstract

The SARS-CoV-2 pandemic has affected more than 70 million people worldwide and resulted in over 1.5 million deaths. A broad deployment of effective immunization campaigns to achieve population immunity at global scale will depend on the biological and logistical attributes of the vaccine. Here, two adeno-associated viral (AAV)-based vaccine candidates demonstrate potent immunogenicity in mouse and nonhuman primates following a single injection. Peak neutralizing antibody titers remain sustained at 5 months and are complemented by functional memory T-cells responses. The AAVrh32.33 capsid of the AAVCOVID vaccine is an engineered AAV to which no relevant pre-existing immunity exists in humans. Moreover, the vaccine is stable at room temperature for at least one month and is produced at high yields using established commercial manufacturing processes in the gene therapy industry. Thus, this methodology holds as a very promising single dose, thermostable vaccine platform well-suited to address emerging pathogens on a global scale.

## INTRODUCTION

A severe acute respiratory disease syndrome caused by a novel coronavirus was first reported in December 2019 (COVID-19 disease) and was subsequently shown to be caused by SARS-CoV-2 (Zhou et al., 2020). A safe and effective vaccine against COVID-19 has been extensively sought since early 2020, when the virus was first isolated and sequenced. A prophylactic vaccination program using an effective vaccine candidate would represent one of the best means to mitigate the health and economic burdens imposed by this pandemic. Based on prior work on SARS-CoV-1 and other respiratory viruses, the SARS-CoV-2 Spike protein (S) was considered an attractive antigen target for the induction of protective immunity to the virus (Folegatti et al., 2020a; Graham et al., 2020; Koch et al., 2020; Martin et al., 2008; Modjarrad et al., 2019; Muthumani et al., 2015; van Doremalen et al., 2020a). This viral envelope glycoprotein engages the ACE-2 cellular receptor through its receptor binding domain (RBD), a key target for developing neutralizing antibodies (Wang et al., 2020). Antigenicity of a full-length S protein can be further enhanced by select proline substitutions that maintain the S protein in a pre-fusion conformation (Walls et al., 2020; Wrapp et al., 2020). The utility of employing this antigen as a vaccination target has been validated by reports of substantial protective efficacy in several human vaccine studies (Folegatti et al., 2020b; Jackson et al., 2020).

The magnitude of this crisis has motivated the initiation of over 200 SARS-CoV-2 vaccines in development across various technology platforms (https://www.who.int/publications/m/item/draft-landscape-of-covid-19-candidate-vaccines). Inactivated viral and gene-based vaccines progressed particularly rapidly with the start of the first Phase 1 studies ocurring about 3 months after the identification and sequencing of SARS-CoV-2. The various gene-based vaccines encode for SARS-CoV-2 or RBD-containing sequences and leverage different gene delivery platforms including unencapsulated or naked DNA delivered by electroporation (Patel et al., 2020; Yu et al., 2020), mRNA delivered mostly by lipid nanoparticles (Anderson et al., 2020; Corbett et al., 2020; Jackson et al., 2020; Kalnin et al., 2020; Kremsner et al., 2020; Walsh et al., 2020; Zhang et al., 2020) and viral vectors such as vesicular stomatitis virus (VSV) (Case et al., 2020), adenovirus (Feng et al., 2020; Folegatti et al., 2020b; Hassan et al., 2020; Logunov et al., 2020; Mercado et al., 2020; van Doremalen et al., 2020b; Zhu et al., 2020a; Zhu et al., 2020b) or yellow fever virus (Sanchez-Felipe et al., 2020). The worldwide COVID-19 vaccine development efforts are in different stages of development, however, less than a year after the start of the outbreak, at least three of the gene-based vaccine candidates appear to be efficacious and have demonstrated favorable short-term safety in large Phase 3 studies and two have received the Emergency Use Approval (EUA) by the FDA (https://www.fda.gov/media/144412/download, https://www.fda.gov/media/144636/download).

As public health programs race to vaccinate individuals with these early wave vaccines, some of the limitations of the first-generation vaccines are becoming increasingly apparent, especially when attributes required for global distribution outside the U.S. and Europe are considered. The cold chain storage requirement and reliance on more than one injection to induce protective immunity are major limitations of some of the first-generation vaccines (Folegatti et al., 2020b; Jackson et al., 2020). It is less clear at this time, but issues such as post-inoculation reactogenicity, durability of immune responses, and efficacy in populations with known vulnerabilities to COVID-19, such as the elderly and obese, may also be attributes upon which second wave vaccines can improve (Biswas et al., 2020; Cuschieri and Grech, 2020). Moreover, little is known at this time about the expense and reliability of scale-up manufacturing that will be critical in assessing the feasibility of using any vaccine in global vaccine campaigns.

Here, we interrogate the safety and immunogenicity of two novel adeno-associated viral (AAV) vector vaccines in mice and nonhuman primates (NHP). In addition, to address some of the logistical and biological hurdles in the global distribution challenge mentioned above, we evaluated whether these genetic vaccines are immunogenic following a single injection, induce immune responses in animal models of obesity and aging, demonstrate long-term durable responses, and retain activity at various storage temperatures.

Wild type AAV is a non-enveloped single stranded DNA dependoparvovirus with broad host range including nonhuman and human primates. AAV serotypes of all characterized versions to date are not associated with human disease, yet many serotypes are endemic and antibodies directed at those serotypes are prevalent in human populations (Calcedo et al., 2009; Gao et al., 2004). Replication defective viral vector particles can be produced by providing in *trans* viral open reading frames to a recombinant genome composed of the transgene and AAV inverted terminal repeats (ITRs), the only viral element retained. Based on studies with natural and engineered variants, the AAV capsid is known to be a primary determinant of biodistribution, immunogenicity, production yield, and efficiency of gene transfer (Salganik et al., 2015; Vandenberghe et al., 2009b).

To date, AAV-based vectors have primarily been used for therapeutic gene transfer in genetically defined diseases such as hemophilia and spinal muscular atrophy type I (High and Roncarolo, 2019). Four decades of AAV research has established an overall favorable safety profile in preclinical and clinical studies and has led to the commercialization of 3 products initially approved by US FDA and European EMA: Glybera, Luxturna, and Zolgensma. AAV is generally well tolerated via several local routes of administration, including intramuscular (IM) injection, and large doses delivered systemically, typically by intravenous injection, have been commonly used in human clinical trials (Bennett et al., 2016; High and Roncarolo, 2019; Mendell et al., 2017; Stroes et al., 2008). Safety, in part, is thought to be due to the low pro-inflammatory responses to the AAV capsid that allows for a tolerogenic or anergic environment to be established toward a self-transgene product (Mingozzi and High, 2017). However, transgenes that are foreign to the host are known to lead to antigen-specific humoral and cellular immunogenicity, tissue inflammation, and the potential elimination of the transgene product or the transgene expressing cell (Gao et al., 2009; Vandenberghe and Wilson, 2007). While undesirable in therapeutic gene therapy applications, the inflammatory potential of AAV has been leveraged in several preclinical and clinical vaccine studies (Lin et al., 2009; Mehendale et al., 2008; Nieto and Salvetti, 2014; Vardas et al., 2010).

AAVrh32.33 is a hybrid serotype capsid that is phylogenetically distinct from commonly used gene therapy AAVs such as AAV2, AAV8 or AAV9, with a sequence identity less than 70% (Figure 1B) (Calcedo et al., 2009; Lin et al., 2009; R et al., 2009; Vandenberghe et al., 2009a). In contrast to most AAVs studied to date, AAVrh32.33, following intramuscular injection (IM), elicits a pro-inflammatory response in mice, declining transgene expression over time, and pronounced local inflammatory infiltrates (Lin et al., 2009; Mays et al., 2009; Mays et al., 2013). Moreover, in an extensive sero-epidemiological study of AAVrh32.33, less than 2% of subjects carried neutralizing responses above a titer of 1:20 (Calcedo et al., 2009). These rare, low titer antibody levels did not reduce vaccine efficacy due to neutralization when AAVrh32.33 was used as the vaccine vector in contrast to what was observed when AAV vectors were used to which higher titer antibodies were prevalent (Lin et al., 2009). Previously, these key attributes of AAVrh32.33 compelled us and others to explore the potential of this AAV serotype as a vaccine platform. This earlier work established viability as potential vaccine via proof-of-concept studies in an influenza murine challenge model, a mouse and NHP HIV immunogenicity study, as well as in applications in dengue and HCV infections (Li et al., 2012; Lin et al., 2009; Zhu et al., 2015; Zhu et al., 2019). Collectively, these studies provided evidence of potent, durable, and functional B and T-cell responses.

**Figure 1.**
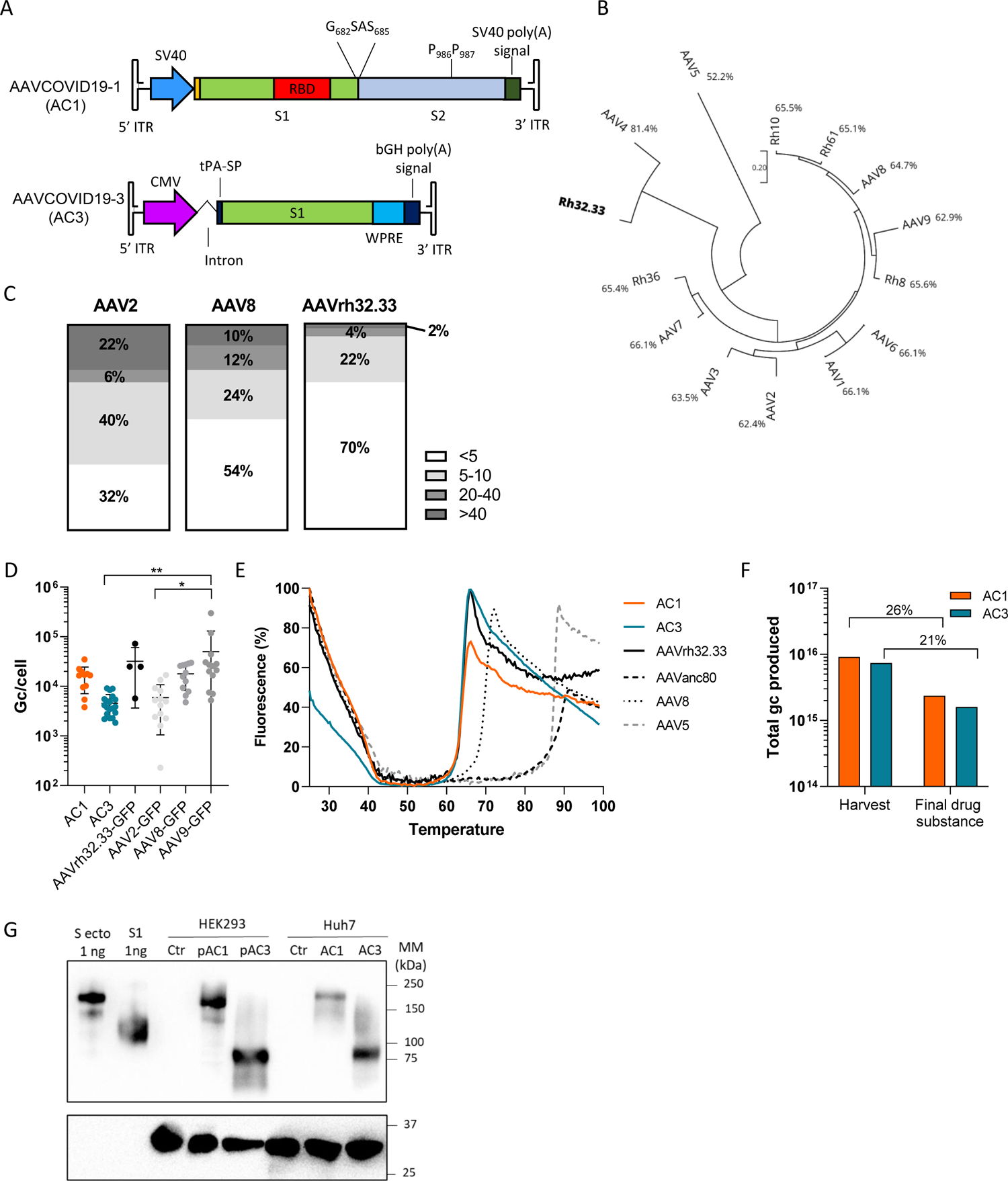
Composition and characterization of AAVCOVID vaccine candidates. (A) Schematic representation of the recombinant genome of AAVCOVID19-1 (AC1) and AAVCOVID19-3 (AC3) vaccine candidates. SV40: Simian virus 40 promoter. RBD: receptor binding domain. S1: SARS-CoV-2 Spike subunit 1. S2: SARS-CoV-2 Spike subunit 2. CMV: cytomegalovirus promoter. tPA-SP: tissue plasminogen activator signal peptide. WPRE: woodchuck hepatitis virus posttranscriptional regulatory element. bGH: bovine growth hormone. ITR: inverted terminal repeat. (B) Phylogenetic tree of several AAV clades and percentage of sequence identity with AAVrh32.33. (C) Percentage of seropositivity of neutralizing antibodies and titer range against AAV2, AAV8 and AAVrh32.33 among 50 donor plasma samples. (D) Productivity of several AC1 and AC3 (vector genome copies produced per producer cell or Gc/cell) compared to various AAV serotypes carrying a CMV-EGFP-WPRE transgene in small scale production and purification. Data are represented as mean ± SD. One-way ANOVA and Tukey’s tests were used to compare groups between them. * p<0.05, ** p<0.01. (E) AAV-ID analysis of capsid identity and stability of AC1 and AC3 compared to AAVrh32.33 and other serotypes. (F) Total genome copies (gc) of AC1 and AC3 produced at large scale and quantified at harvest of producer cells and after the purification (final drug substance). Percentage of vector recovery during purification is displayed. (F) Detection of SARS-CoV-2 Spike antigens by Western blot in HEK293 cells transfected with 1 µg of ITR-containing pAC1 or pAC3 plasmids and Huh7 cells transduced with 5×10^5^ gc/cell of AC1 and AC3 72h after treatment. Recombinant S ectodomain (S ecto, lane 1) and S1 subunit (S1, His-tagged, lane 2) were used as positive control and size reference.

Upon the news of the emerging epidemic in Wuhan, China and the sequencing of the likely etiological agent (Wu et al., 2020), we initiated the development of several AAVrh32.33-based vaccine candidates (AAVCOVID). Two gene-based vaccine candidates were selected, which encode different versions of the SARS-CoV-2 S antigen. AAVCOVID19-1 (AC1) encodes a full-length S protein locked in a prefusion conformation that is designed to be expressed on the cell membrane of transduced cells, whereas AAVCOVID19-3 (AC3) candidate’s S protein is shorter and engineered to be secreted.

Here, the two AAVCOVID vaccine candidates are shown to elicit strong B and T cell immunogenic responses following a single intramuscular injection in rodents and NHPs. These immunogenic responses have proven long-lived, with levels remaining at peak or near peak levels for at least 5 months following the initial injection. AAVCOVID is minimally reactogenic in NHPs, can be manufactured at scale with standard industrial processes, and importantly, remains stable during storage at room temperature for at least one month. Combined, these attributes make AAVCOVID a promising candidate for a safe, effective and protective vaccine, amenable to manufacturing at scale and broad distribution that could contribute to the challenge of addressing the global need for vaccines against COVID-19.

## RESULTS

### Design and production of AAVCOVID vaccines

AC1 and AC3 are both viral vector COVID-19 vaccine candidates composed of an AAVrh32.33 capsid and an AAV2 ITR-flanked transgene expressing distinct SARS-CoV-2 S antigens. Figure 1A depicts AC1 which encodes a full-length membrane anchored S protein based on the Wuhan sequence, modified by amino-acid substitutions that prevent S1/S2 furin cleavage and stabilize S in a pre-fusion conformation for optimal RBD exposure and antigenicity (Walls et al., 2020; Wrapp et al., 2020). AC3 expresses the secreted S1 subunit of the Wuhan S protein (Figure 1A). AAVrh32.33 is a previously described rhesus derived AAV serotype. It is most closely related to AAV4 but phylogenetically divergent from the AAVs that are most commonly circulating and used as gene therapy vectors in humans (Figure 1B). Previously it was shown in an extensive human epidemiological study that the seroprevalence of antibodies to AAVrh32.33 is minimal (Calcedo et al., 2009). Consistent with these findings, 50 plasma samples collected from healthy donors demonstrated highly reduced antibody prevalence to AAVrh32.33 as compared with that seen to AAV8 and AAV2, with 6% of samples with titers of 1:20 or above compared to 22% and 28% respectively (Figure 1C). In terms of production yields, in more than 10 research grade preparations, AC1 was shown to be comparable to serotypes AAV8 and AAV9, while AC3 showed slightly reduced productivity (Figure 1D). The capsid identity of AC1 and AC3 is consistent with AAVrh32.33 in the AAV-ID thermostability assay (Figure 1E) (Pacouret et al., 2017).

Given the need for scaled production of vaccines, we evaluated whether AC1 and AC3 manufacturing was feasible with a previously established scalable process developed by Novartis Gene Therapies. Specifically, approximately 1.5 x 10^10^ HEK293 were seeded in a fixed-bed bioreactor (PALL iCellis 500), grown for 4 days, and then co-transfected with the AAVCOVID ITR plasmids, pKan2/rh32.33 for the AAV capsid and pALDX80 as helper. At harvest, yields for AC1 and AC3 were above 7 x 10^15^ genome containing particles or genome copies (gc). Subsequent purification included multiple tangential flow filtration (TFF), an ion-exchange chromatography and a cesium chloride density gradient step to eventually recover between 21% and 26% in the final drug substance (Figure 1F). Of note, given the expedited nature of these studies, less than 2 weeks of process development studies were performed to successfully adapt the existing scalable production system to AAVrh32.33.

Lastly, expression of the S transgene was detected for each AAVCOVID candidate *in vitro* by transfection and transduction (Figures 1G and S1). Higher expression of AC3 was detected at mRNA (Figure S1A) and protein level (Figures 1G and S1).

### A single dose of AAVCOVID induces high and durable antibody titer in two mouse strains

The immunogenicity of AC1 and AC3 following a single injection at a low and high dose of 10^10^ and 10^11^ gc, respectively, in the gastrocnemius muscle was evaluated in 6-10-week-old BALB/C and C57BL/6 mice of both genders. SARS-CoV-2 (SARS2) RBD-binding IgG antibody levels were monitored by ELISA at regular intervals (Figures 2A and 2B), as were neutralizing antibody levels assayed using a SARS-CoV-2 Spike pseudotyped lentivirus (pseudovirus) inhibition-of-transduction method (Figures 2C and 2D).

**Figure 2.**
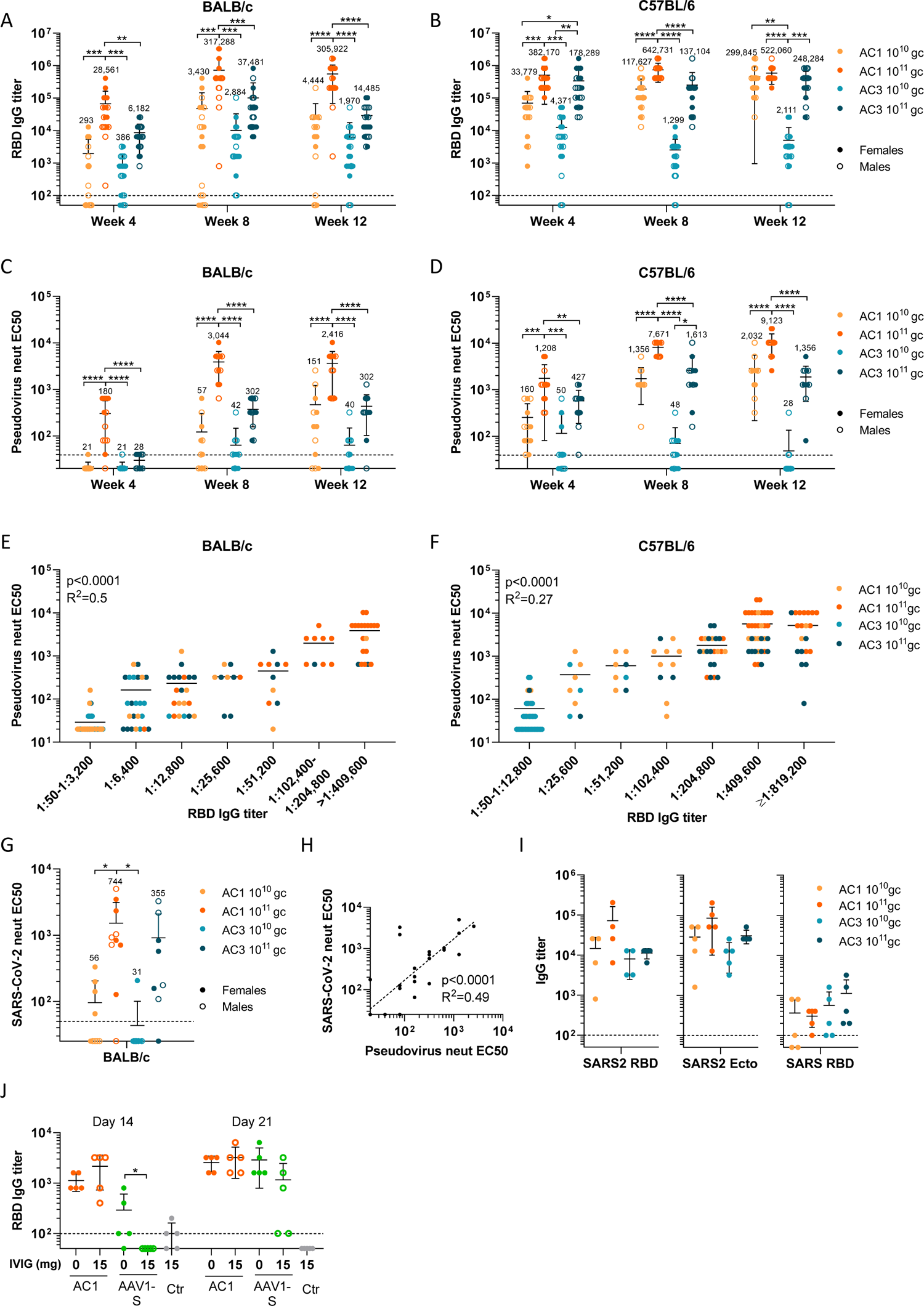
Quantitative assessment of humoral responses in two mouse strains. (A-B) Monthly monitoring of SARS-CoV-2 RBD-binding IgG titers in 6-10 week-old BALB/c (A) and C57BL/6 (B) mice injected IM with two doses (10^10^ gc and 10^11^ gc) of AC1 or AC3, n=20 (10 females and 10 males). Mean geometric titers (MGT) shown above each group. (C-D) Pseudovirus neutralizing titers of a subset of BALB/c (C) and C57BL/6 (D) animals (6 females and 6 males per group) from the studies described in A and B. TheGMT are shown above each group. (E-F) Correlation of pseudovirus neutralizing titers and RBD-binding IgG titers in BALB/c (E) and C57BL/6 (F). (G) Live SARS-CoV-2 neutralizing titers measured on a PRNT assay on week 4 samples harvested from BALB/c animals (n≥8, both genders). The GMT is shown above each group. (H) Correlation of SARS-CoV-2 neutralizing and pseudovirus neutralizing titers. (I) Titer of binding antibodies against SARS-CoV-2 RBD (SARS2 RBD), SARS-CoV-2 Spike ectodomain (SARS2 Ecto) and SARS-CoV RBD (SARS RBD) in female BALB/c sera 28 days after AC1 or AC3 injection. (J) RBD-binding antibody titers in BALB/c male animals (n=5) vaccinated with 10^11^ gc of AC1 or AAV1-S (same genomic sequence packaged in different capsids), which were naïve (0 mg IVIG) or passively pre-immunized with 15 mg of human IVIG 24h and 2h prior to the vaccination. Ctr: unvaccinated control. (A-J) Data are represented as mean ± SD. For (A-D and G) groups were compared by one-way ANOVA and Tukey’s post-test. * p<0.05, ** p<0.01, *** p<0.001, **** p<0.0001. For (E, F, H) Pearson’s correlation coefficient was calculated to assess correlation. For (J) naïve and immunized groups were compared by Mann-Whitney’s U test.

Both mouse strains demonstrated dose-dependent potent binding and neutralizing responses from a single dose administration of AC1 or AC3 that persisted through 3 months. Overall, AC1 at high doses induced a significantly higher level of binding and neutralizing antibody titers to SARS-CoV-2 (binding geometric mean titer (GMT) of 305,922 and 522,060 in BALB/c and C57BL/6, respectively; and neutralizing GMT of 2,416 and 9,123, 12 weeks post-vaccination) than AC3 (binding GMT of 14,485 and 248,284 in BALB/c and C57BL/6, respectively; and neutralizing GMT of 302 and 1,356, 12 weeks post-vaccination). At a low dose, AC1 was superior to AC3, particularly in C57BL/6 mice at later timepoints (Figures 2A-2F). Immunogenicity was modestly lower in males versus female mice for both candidates (Figures 2A-2D). The kinetics of binding-antibody induction showed early onset of responses by day 14 (Figure S2A) and increasing seroconversion rates over time (Figure S2B). Neutralizing antibody kinetics lagged by approximately a week, with limited seroconversion at week 4 that increased thereafter (Figures 2C and 2D). Binding and neutralizing titers correlated; however, AC1 achieved higher neutralizing titers and a larger relative ratio of neutralizing to binding titers compared to those produced by AC3 (Figures 2E and 2F).

Limited plaque reduction neutralizing assay titers (PRNT) with live SARS-CoV-2 were obtained for AC1 and AC3 in BALB/c mice 4 weeks after vaccination, showing the quality of response in terms of the neutralization of SARS-CoV-2 live virus (Figure 2G). These responses correlated modestly well with results from the pseudovirus neutralization assay (Figure 2H). ELISA IgG titers to SARS-CoV-2 S full length ectodomain (SARS2 Ecto), SARS-CoV-1 S RBD (SARS RBD) or MERS S RBD were assayed (Figure 2I). Antibody responses to full length S ectodomain (SARS2 Ecto) were modestly higher compared to RBD titers (SARS2 RBD) (Figure 2I). Cross-reactivity of the elicited IgG with SARS RBD was noted, but at reduced levels (Figure 2I), with no cross-reactivity detected against MERS RBD (data not shown).

Lastly, to model the impact of AAV capsid pre-existing immunity on AAVCOVID immunogenicity in humans, 24 and 2 hours before vaccination BALB/c mice received 15 mg of intravenous immunoglobulin (IVIG) derived from pooled samples from thousands of human donors. As a control, a single dose immunization using the AC1 vector was compared to vaccination with an AAV1 capsid vector containing an identical genome (AAV1-S). AAV1 is known to have higher pre-existing immunity in human populations (Calcedo et al., 2009). Figure 2J shows that animals vaccinated with AC1 were unaffected by the IVIG pretreatment, while AAV1-S had reduced seroconversion on day 21 compared to IVIG-naïve animals.

### AC1 elicits qualitatively distinct humoral response compared to AC3

Next, we assessed the quality of the humoral responses over time for each of the vaccine candidates in BALB/c mice. In AC1-treated animals, IgM and IgA antibodies directed at SARS2 RBD were detected at early timepoints, day 7 and 14 respectively, but the IgG isotype dominated circulating SARS2 RBD-specific antibody levels from thereon (Figure 3A). AC3 IgM and IgA levels were lower than those observed for AC1. Total IgG levels were composed of all IgG subclasses when AC1 was utilized, whereas IgG1 predominated when AC3 was injected. The ratio of IgG2a/IgG1 suggests a balanced Th1 response stimulated by AC1, with more Th2 skewing seen in the AC3 response (Figures 3A and 3B).

**Figure 3.**
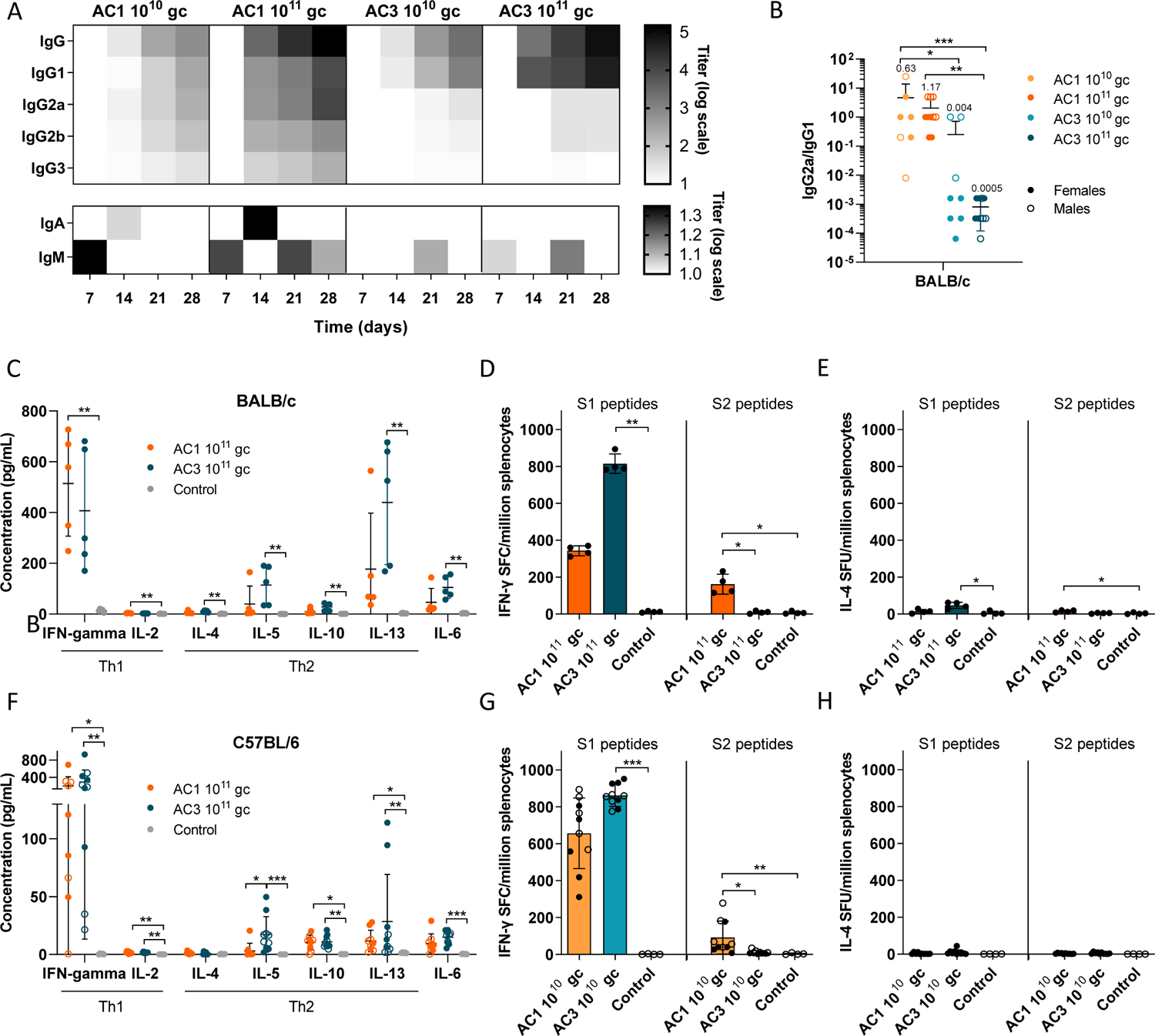
Quality of the host response to AAVCOVID. (A) Several RBD-binding antibody isotype titers (IgG, IgG1, IgG2a, IgG2b, IgG3, IgA and IgM) measured weekly in 6-10 week-old BALB/c (n=10, 5 females and 5 males) treated IM with two doses of AC1 and AC3. (B) Ratio of RBD-binding IgG2a and IgG1 antibody titers in serum samples harvested 28 days after vaccination of BALB/c mice as described in A. The Geometric Mean Titer (GMT) is shown above each group. (C and F) Cytokine concentration (pg/mL) in supernatants harvested from splenocytes stimulated for 48h with peptides spanning SARS-CoV-2 Spike protein. Splenocytes were extracted from BALB/c (C) and C57BL/6 (F) animals 4 and 6 weeks, respectively, after vaccination with 10^11^ gc of AC1 or AC3. (D-E) Spot forming units (SFU) detected by IFN-γ (D) or IL-4 (E) ELISpot in splenocytes extracted from BALB/c animals 4 weeks after vaccination with 10^11^ gc of AC1 or AC3 and stimulated with peptides spanning SARS-CoV-2 Spike protein for 48h. (G-H) Spot forming units (SFU) detected by IFN-γ (G) or IL-4 (H) ELISpot in splenocytes extracted from C57BL/6 animals 6 weeks after vaccination with 10^10^ gc of AC1 or AC3 and stimulated with peptides spanning SARS-CoV-2 Spike protein for 48h. For (B-H) data are represented as mean ± SD and groups were compared by Kruskal Wallis and Dunn’s post-test.

To further interrogate this divergent qualitative response, cytokine secretion and ELISPOT analyses were performed on splenocytes from AC1 and AC3 immunized BALB/c and C57BL/6 animals. Secretion of several cytokines was detected in stimulated splenocytes (Figures 3C and 3F). However, IFN-γ was predominantly secreted and minimal levels of Th2-associated cytokines, such as IL-5 and IL-13, were measured, except in BALB/c mice, where AC3 induced a greater IL-13 response (Figures 3C and 3F).

IFN-γ ELISPOT revealed a robust response against peptides spanning the S1 subunit (Figures 3D, 3G, S3A and S3C), while lower responses were detected against the S2 subunit only in the AC1 vaccinated group. Minimal IL4 responses were seen by IL4 ELISPOT (Figures 3E, 3H, S3B and S3D).

### Immunogenicity of AAVCOVID is influenced by age but retains potency in obese mice

Vaccine efficacy is often impaired in obese or elderly humans, which are two of the most vulnerable populations in the COVID-19 pandemic. To model this conditions, 18-week and 2-year-old mice of both genders were immunized with AAVCOVID at low and high doses, bled at regular intervals, and analyzed for SARS2 RBD IgG and pseudovirus neutralization responses in the serum. A reduction in IgG and neutralizing titers is observed between 18-week and 2-year-old mice (Figures 4A-4D). 18-week-old and, to a lesser extent, 2-year-old mice developed robust neutralizing titers upon vaccination with AC1 (Figures 4B and 4D), but the AC3 at high doses failed to recapitulate the results in younger mice (Figure 2D). Low doses and high dose of AC3 failed to elicit neutralizing antibodies in most of the 2-year-old mice, while animals treated with a high dose of AC1 showed high titers but incomplete seroconversion (Figures 4D and 4E). In aggregate, high dose AC1 demonstrates robust, albeit reduced immunogenicity in aged mice, with clearly superior immunogenicity compared to that induced by AC3 in aged mice.

**Figure 4.**
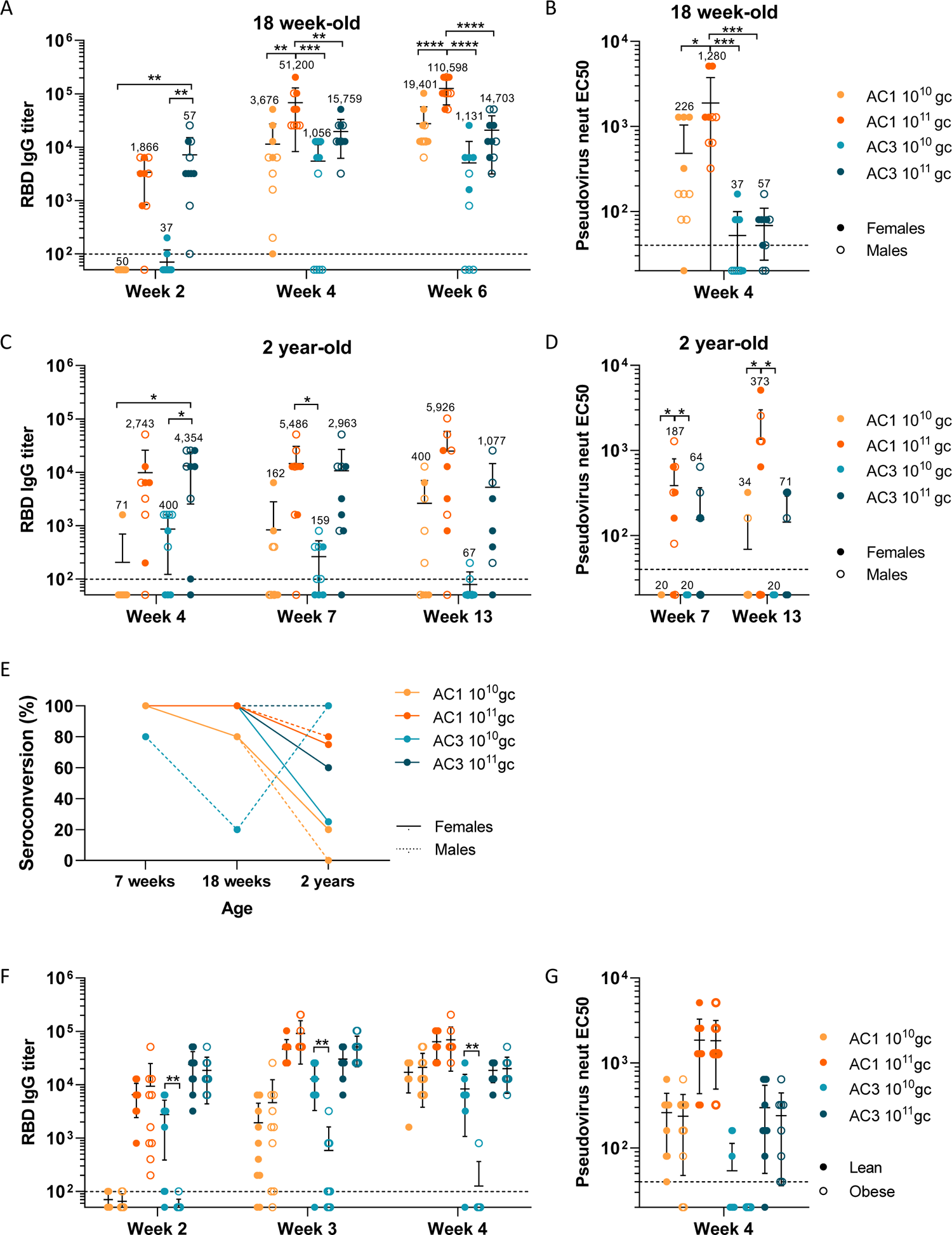
Humoral responses in murine models of age and obesity. (A) RBD-binding antibody titers measured on weeks 2, 4 and 6 in 18 week-old C57BL/6 animals (n≥9, both genders) vaccinated with two doses (10^10^ gc and 10^11^ gc) of AC1 and AC3 intramuscularly. Mean geometric titers (MGT) shown above each group. (B) Pseudovirus neutralizing titers on week 4 in animals described in A. The Geometric Mean Titer (GMT) is shown above each group. (C) RBD-binding antibody titers measured on weeks 4, 7 and 13 in 2 year-old C57BL/6 animals (n≥7, both genders) vaccinated with two doses (10^10^ gc and 10^11^ gc) of AC1 and AC3 intramuscularly. GMT is shown above each group. (D) Pseudovirus neutralizing titers on weeks 7 and 13 in animals described in C. GMT is shown above each group. (E) Seroconversion rates in RBD-binding antibodies 4 weeks after vaccination of C57BL/6 mice at different ages. (F) RBD-binding antibody titers measured on weeks 2, 3 and 4 in 12-week-old lean and obese C57BL/6 animals (n=10 males) vaccinated with two doses (10^10^ gc and 10^11^ gc) of AC1 and AC3 intramuscularly. (G) Pseudovirus neutralizing titers on week 4 in animals described in F. (A-D, F-G) Data are represented as mean ± SD. For (A-D) groups were compared by one-way ANOVA and Tukey’s post-test. (F-G) Lean and obese mice receiving the same treatment were compared by Student’s t test. * p<0.05, ** p<0.01, *** p<0.001, **** p<0.0001.

A diet-induced C57BL/6 obesity (DIO) mouse model was used to study vaccine efficacy in inducing SARS2 RBD-specific antibodies in overweight animals. 12-week-old C57BL/6 and C57BL/6 DIO (n=10) mice were vaccinated with 10^10^ and 10^11^ gc of AC1 and AC3. IgG RBD-binding and neutralizing antibody levels were indistinguishable between lean and obese groups for AC1 and the high dose group of AC3, yet interestingly the low dose of AC3 produced a less robust antibody response in the DIO mice than did the comparable dose of AC1 (Figures 4F and 4G).

### Durable neutralizing antigenicity in NHP from a single dose injection

To model the immunogenicity of AAVCOVID in humans, one female and one male rhesus macaque were injected IM with 10^12^ gc of AC1 and AC3. Animals tolerated the vaccine dose well, with no temperature elevations or local reactogenicity based on clinical examinations, complete blood counts and chemistry (Figure S4), or cytokine analysis (Figure S5). Regular phlebotomies were performed to assess RBD binding, pseudovirus neutralizing, and live SARS-CoV-2 neutralizing antibody titers in serum (Figures 5A, 5B and 5C) and B cell analysis from PBMCs (Figures 5D, 5E and 5F). These animals continue to be monitored to assess the durability of the vaccine response and are currently at the 5-month time point following the single dose immunization.

**Figure 5.**
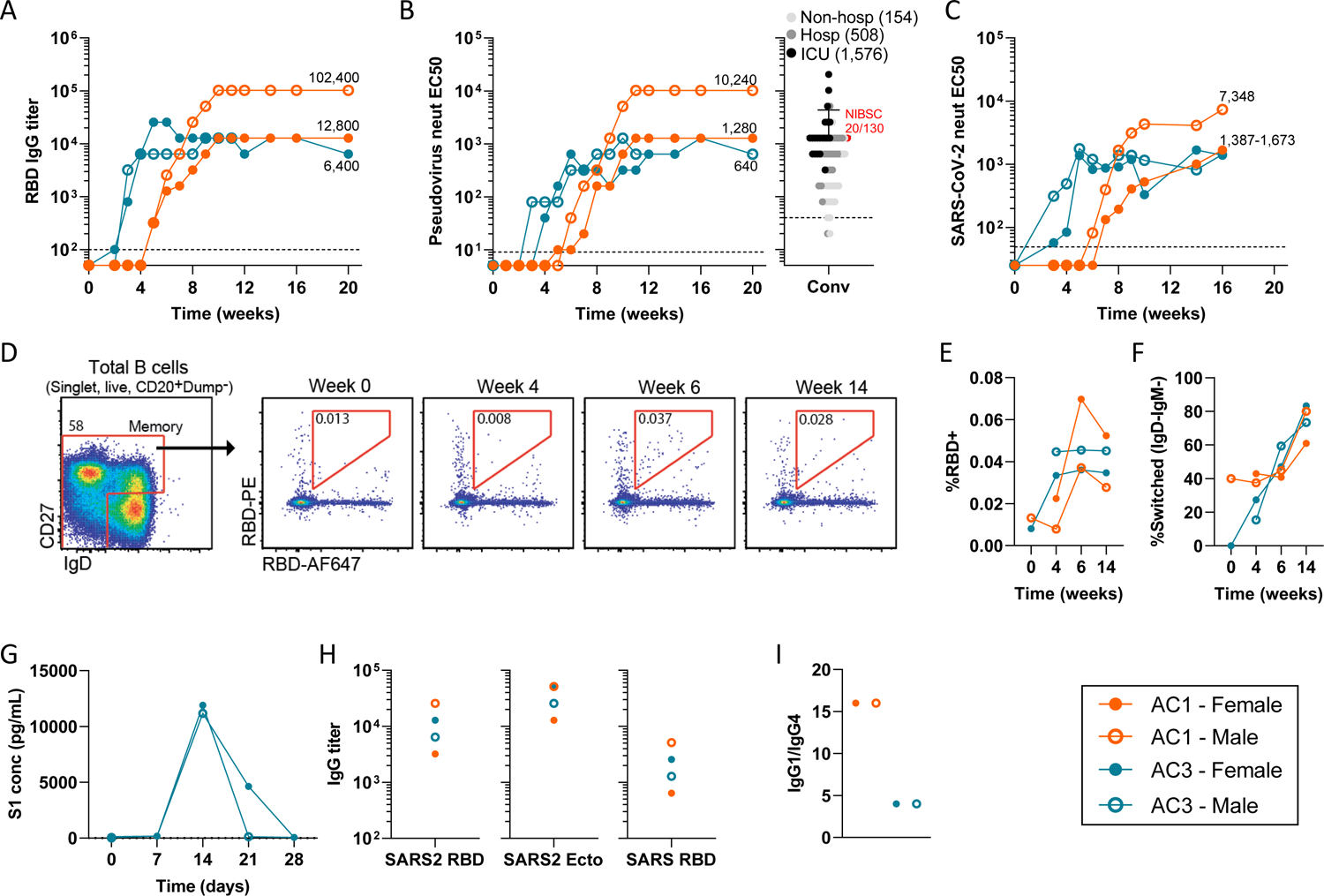
Characterization of humoral immune responses in NHP. (A) SARS-CoV-2 RBD-binding IgG titers 20-week follow up in Rhesus macaques (n=2, 1 female and 1 male) treated IM with 10^12^ gc of AC1 or AC3. (B) Pseudovirus neutralizing antibody titers in NHPs for 20 weeks (left) and 60 convalescent human plasma samples of patients with different disease severity and NIBSC 20/130 (red dot) reference plasma (right). The Geometric Mean Titer (GMT) is shown for each cohort of convalescent plasma. (C) Live SARS-CoV-2 neutralizing titers. (D) Identification of RBD-binding B cells with a memory phenotype (CD27+ or CD27-IgD-) in peripheral blood of a representative macaque at multiple dates post-vaccination. (E) Frequency of RBD-binding B cells in memory B cell compartment. (F) Frequency of RBD-binding memory B cells with isotype-switched (IgD-IgM-) phenotype. (G) Quantification of S1 subunit concentration (pg/mL) in sera of animals treated with AC3 during the first month after vaccination. (H) Titration of binding antibodies against SARS-CoV-2 RBD (SARS2 RBD), SARS-CoV-2 Spike ectodomain (SARS2 Ecto) and SARS-CoV RBD (SARS RBD) 9 weeks after vaccination. (I) Ratio between RBD-binding IgG1 and IgG4 isotypes 8 weeks post-vaccination.

AC3 SARS2 RBD-binding antibody responses were detectable as early as week 3 after a single administration and plateaued by week 5 hovering around 1:6,400 and 1:12,800 (Figure 5A). AC1 IgG, on the contrary, only became apparent on week 5 and then steadily increased until week 10. One AC1-injected animal achieved similar antibody levels to those measured in both AC3 vaccinated primates (1:12,800) while the other AC-1 vaccinated animal achieved levels that were 8-fold higher (Figure 5A). With minimal fluctuation, SARS2 RBD IgG levels have been maintained to date at peak levels, now 20 weeks or 5 months after a single shot vaccine for both the AC1 and AC3 injections.

Pseudovirus neutralizing titers and PRNT closely tracked with a slight delay in the IgG kinetics for both AC1 and AC3, reaching peak neutralizing titers 6 to 8 weeks after vaccination for AC3 (1:640 and 1:1,280) and 11 weeks following AC1 injection (1:1,280 and 1:10,240). These neutralizing antibody responses have remained stable at peak levels through week 20 in the pseudovirus neutralizing assay and 16 weeks in the PRNT assay (Figures 5B and 5C), the last time points analyzed. Benchmarking of the pseudovirus neutralizing assay was performed in 2 ways. First, 60 human convalescent plasma samples from 3 cohorts were analyzed (Figure 5B) (non-hospitalized (GMT: 154), hospitalized yet not critical (GMT: 508) and ICU patients (GMT: 1,576)), which demonstrated a clear increase of neutralizing titers with severity of disease. Second, a provisional World Health Organization recommended reference plasma (NISBC 20/130) yielded a 1:1,280 titer in our assay, which was in line with the reported values (Figure 5B). In summary, AC1 induces neutralizing titers in the range of 1:1,280 and 1:10,240 which is in the higher range of convalescence of hospitalized and ICU patients while AC3 leads to titers of 1:640-1:1,280 which is in the range of hospitalized non-ICU patients. These titers persist for at least 5 months.

To track vaccine-induced peripheral blood B cells, a double-labeling technique with fluorophore-conjugated SARS2 recombinant RBD protein was utilized (Figure 5D) (Johnson et al., 2020; Knox et al., 2017). RBD-binding memory B cells (MBCs) were absent at week 0 and detectable by week 4 in three of the animals (Figure 5E). RBD-specific MBCs peaked in frequency at 6 weeks post-vaccination in all recipients and were maintained at a similar level at least through week 14 (Figure 5E). Surface immunoglobulin isotype analyses found an early bias toward generation of IgM-expressing MBCs, whereas isotype switched (IgD^-^IgM^-^; likely IgG^+^) MBCs dominated the SARS2 RBD-specific response by week 14 (Figure 5F). These findings suggest durable induction of SARS2 RBD-specific memory B cells by both AC1 and AC3 vaccines.

Interestingly, in AC3 injected primates, the secreted S1 protein was detectable in their serum 2 weeks after injection. However, the S protein returned to undetectable levels in both animals by week 4, concurrent with increasing anti-SARS2 RBD antibody titers (Figure 5G). Similar to the mouse data, SARS2 ectodomain IgG levels in NHP were higher than SARS2 RBD IgG, and modest cross-reactivity to SARS1 RBD was detected (Figure 5H). Total IgG for both AC1 and AC3 animals was primarily composed of IgG1, suggestive that in NHP, as opposed to mice, both responses appear more Th1-like (Figure 5I).

### Memory T cell response to Spike antigen is developed in NHP

T cell responses to transgene peptide pools (Figure S6A) were analyzed by IFN-γ ELISPOT (Figures 6A and 6B) and intracellular cytokine staining (ICS) (Figures 6C-6F) from PBMCs harvested at monthly intervals. AC3 injected animals showed responses specific to the S1 subunit, higher in the female, starting on week 4 (Figure 6B); however, lower responses were detected in the AC1 female starting on week 8 and there was only a minimal response in the AC1 vaccinated male (Figure 6A).

**Figure 6.**
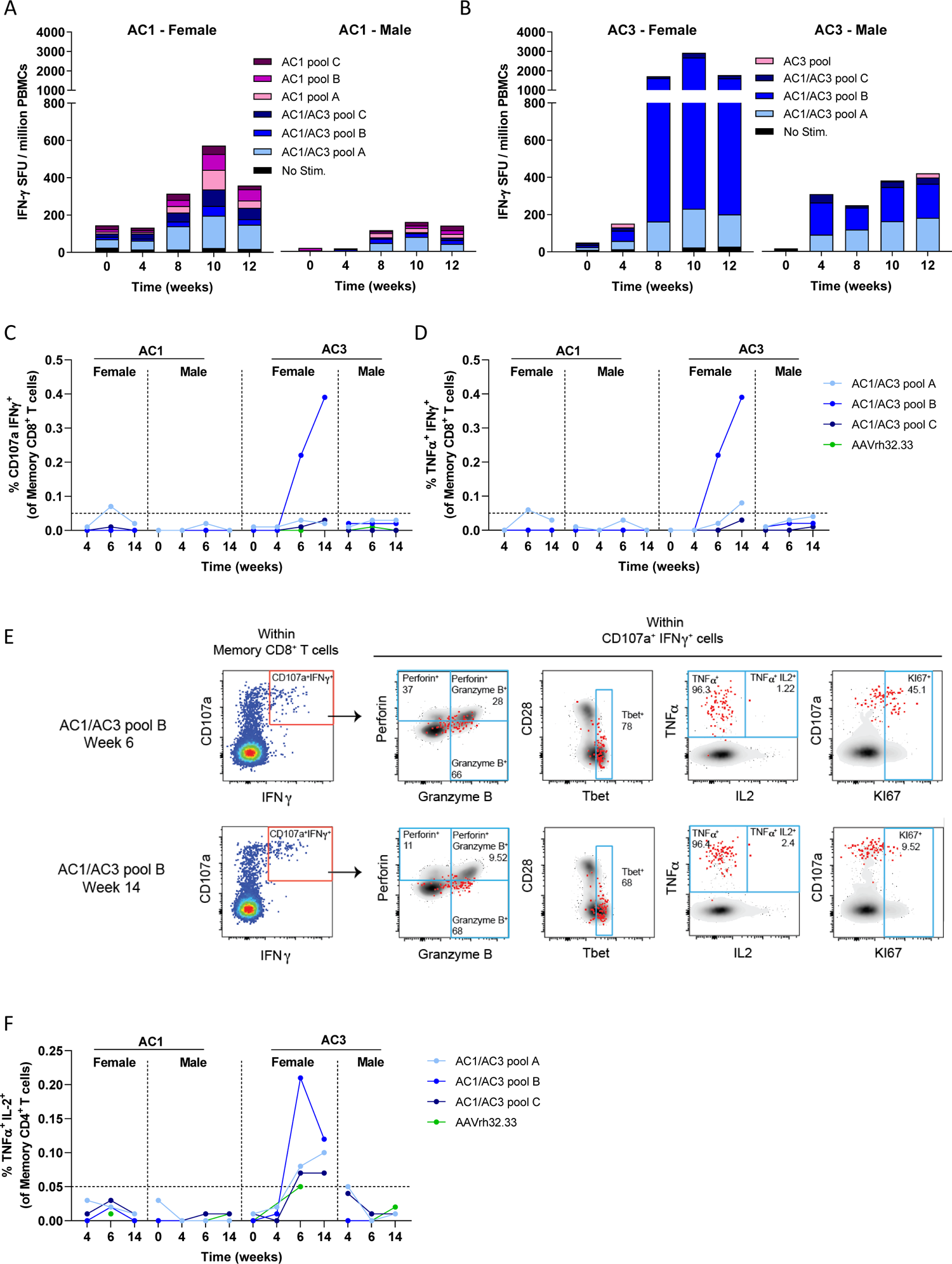
Characterization of cellular immune responses in NHP. (A-B) Quantification of spot forming units (SFU) by ELISpot in PBMC samples collected at different timepoints in rhesus macaques (n=2/vector) treated with AC1 (A) or AC3 (B) and stimulated with peptides specific for each transgene. (C-D) Dot plots summarizing the background subtracted frequency of CD107a^+^IFNγ^+^ or TNFα^+^ IFNγ^+^ cells responding to AC1/AC3 and AAVrh32.33 peptide pools at baseline and at different time points after vaccination. The dotted line indicates the cutoff for positive responses. (E) Flow cytometry plots from AC3 female indicating the frequency of Perforin, Granzyme B, Tbet, TNFα, IL2 and KI67-positive cells within CD107^+^ IFNγ^+^ Memory CD8+ T cells responding to AC1/AC3 shared peptide pool B at day 42 and 98 post vaccination. In the flow plots, total CD107^+^ IFNγ^+^ cells were depicted as red dots overlayed on total Memory CD8^+^ T cells (black). (F) Dot plots summarizing the frequency of background subtracted TNFα^+^ IL2^+^ cells responding to AC1/AC3 and AAVrh32.33 peptide pools at baseline and at different time points after vaccination (n=4). The dotted line indicates the cutoff for positive responses.

Flow cytometry was used to identify the phenotype and functionality of S-specific cells after stimulating PBMCs with the overlapping S1 peptides (note that S2-specific responses in AC1 animals, which were clearly detected by ELISPOT, were not studied in this analyses). The female AC3 showed a robust memory CD8^+^ T cell response to the S1 subunit beginning at week 6 (Figures 6C, 6D and S6B). CD107a^+^ IFNγ^+^ responding cells also produced TNFα at week 6 and 14 (Figures 6D and 6E), but the cytotoxic and activation profile changed over time. At week 6, the S1 subunit-specific cells showed higher expression of the cytotoxicity markers perforin and granzyme B and the activation marker KI67, compared to week 14 (Figure 6E). S1-specific memory CD4^+^ T cell responses were also detected through production of TNFα and IL2 in the female treated with AC3 at week 6 and 14, although these were proportionately lower compared to the corresponding memory CD8^+^ T cell responses (Figures 6F and S6C).

### NHPs develop slow neutralizing antibody response to AAVrh32.33 capsid that shows no cross-reactivity with other AAV serotypes

Viral vectored vaccines are known to induce responses to the delivery vector component, in this case to the AAV capsid. These can enhance the overall immunogenicity of the vaccine, influence its reactogenicity, or prevent the effectiveness of subsequent dosing with a homologous vector due to the neutralization of the vector upon re-administration (Greig et al., 2016; Majowicz et al., 2017). Similarly, in the context of AAV, the cross-reactivity of these antibodies may affect subsequent applications of alternative AAV serotypes that could be neutralized via cross-reactive antibodies to AAVrh32.33, thus potentially influencing future applications of gene therapy for subjects vaccinated with AAVCOVID. In this rhesus study, Table S1 shows that AAVrh32.33 neutralizing antibodies did develop, albeit with slow kinetics and to relatively low levels. Importantly, these modest AAV neutralizing responses did not exhibit cross-neutralization of a panel of commonly used AAV gene therapy serotypes AAV1, 2, 5, 8, and 9 (Table S1 and Figure S7A). In addition, no significant increase in cellular responses against capsid peptides were detected in PBMCs up to 2 months after vaccination (Figure S7B).

### Vector is retained in the injection site and cleared over time in mouse

A biodistribution of the vector following AAVCOVID intramuscular injection was analyzed to establish the kinetics of transgene expression and identify which tissues were transduced beyond that of the intended muscle target (Figure S8). Previously, an AAVrh32.33 expressing a non-self-transgene, when injected intramuscularly in mice, showed declining transgene expression over time that was associated with increasing inflammatory infiltrates at the injection site several weeks after injection (Mays et al., 2009; Mays et al., 2013). This is in stark contrast to other AAVs expressing the same transgene which led to stable transgene expression and minimal local inflammation (Mays et al., 2009; Mays et al., 2013). In the current experiment, C57BL/6 mice were injected with 10^11^ gc in the right gastrocnemius muscle.

Animals were euthanized 1, 4 and 8 weeks after vaccination and tissues were analyzed for vector genome copies and transgene expression. As observed in Figure S8A, vector genome copies in the injected muscle decreased more than 20-fold from week 1 to week 8. AC3 transgene expression declined in a manner similar to the decline in DNA vector genome copy number. Remarkably, AC1 transgene expression was lower than AC3 expression, close to background levels, possibly due to lower promoter activity (Figure S8B). Gene transfer and transduction levels of the contralateral gastrocnemius muscle, liver, and spleen demonstrated 10-100-fold less vector DNA at week 1 than measured in the injected muscle with a steady decline of vector DNA and RNA, at times to undetectable levels (Figures S8A and S8C). A more comprehensive biodistribution study in BALB/c mice that received the 10^11^ gc IM dose of AC3 and were euthanized at week 8 further indicated that the predominant tissue of vector genome and transgene expression was the injected muscle (Figure S8D).

### AAVCOVID is stable and retains potency after one-month room temperature storage

To interrogate the cold chain requirements for storage and transportation of AAVCOVID, research grade vaccine preparations were aliquoted and stored at different temperature conditions (−80°C, 4°C or room temperature (RT)) for 1, 3, 7 or 28 days. Physical vector stability was assessed by titration of DNAse resistant vector genomes and loss or degradation was assessed by comparison to vector aliquots stored at − 80⁰ C (Figure 7A and Table S2). After being stored at 4°C or RT, neither AC1 nor AC3 show a reduction of titers for at least one month. In addition, potency was assessed by injection of 5 x 10^10^ gc of AC1 aliquots in female BALB/c mice. Animals vaccinated with AC1 stored at 4°C or RT for up to 28 days showed similar levels of antibody compared to a control group that received vaccine vectors stored at −80°C (Figure 7B). Although not significant, antibody titers trended downwards with time. Larger studies need to be performed to elucidate if potency can be maintained for longer periods.

**Figure 7.**
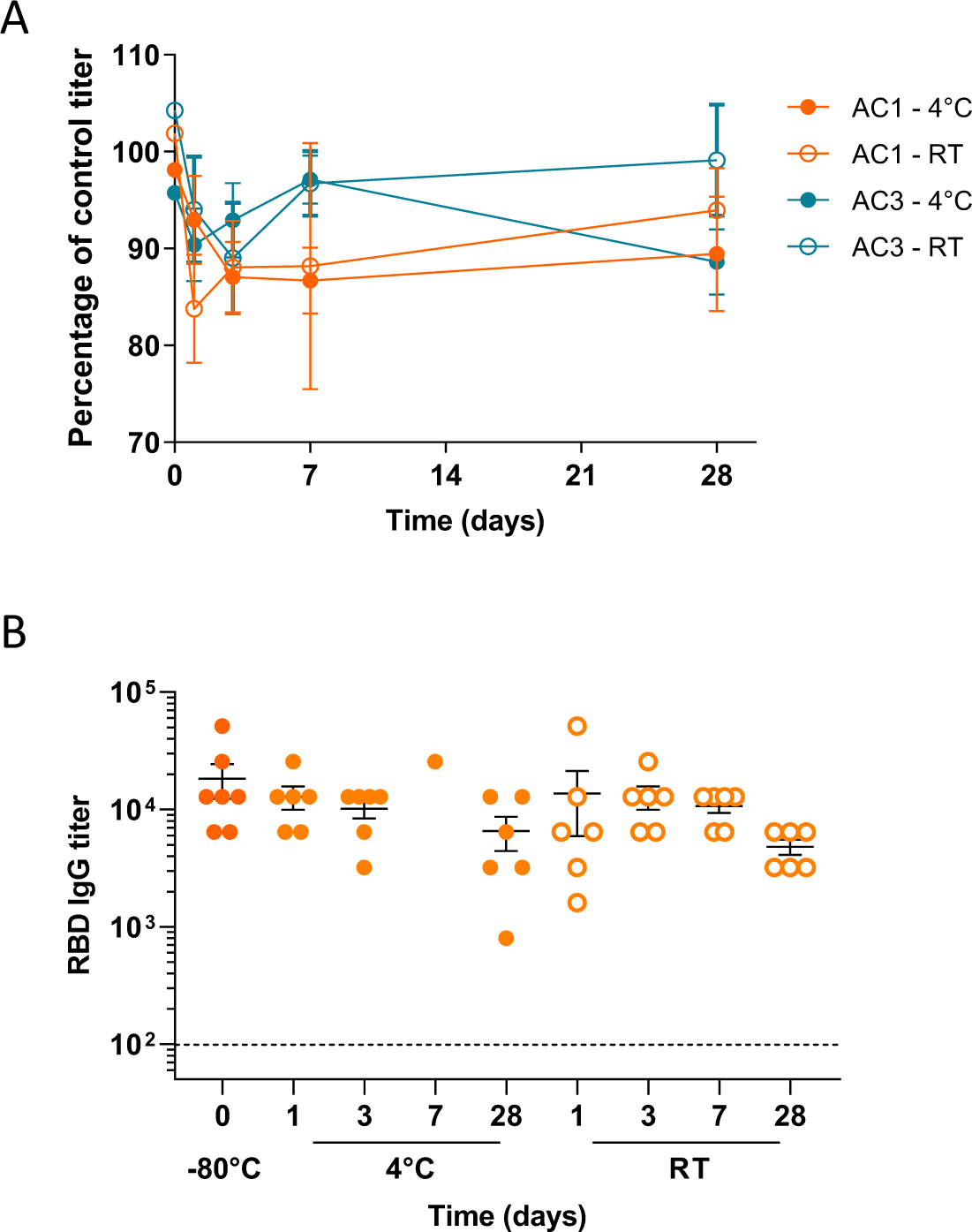
AAVCOVID stability under various storage temperatures. (A) Percentage of titer relative to the −80°C stored control for AC1 and AC3 aliquots stored at 4°C or room temperature (RT) for 1, 3, 7 or 28 days. (B) Measurement of RBD-binding IgG titers in BALB/c female animals vaccinated with AC1 aliquots kept at several temperatures for 1,3, 7 and 28 days. Animals received 5×10^10^ gc IM and antibodies were measured 24 days post-vaccination.

## DISCUSSION

The development of safe and effective vaccines is vital to the worldwide effort to reduce the burden on global health and economic vitality that has been disrupted by the SARS-CoV-2 pandemic. An unprecedented and remarkable effort has led several vaccine candidates to be authorized or approved for commercial use in less than a year since the etiological agent was identified, and several others are nearing that milestone. However, beyond safety and efficacy, other vaccines attributes will likely be critical to achieving the desired long-lasting herd immunity at a global population scale. Many of these considerations are logistical in nature and seek to reduce the cost, time, and complexity of vaccine distribution using available infrastructure around the world. These considerations are particularly germane to producing effective vaccine campaigns in developing nations. Biological features, such as potency from a single dose, durability of protection, and stability at ambient temperature, are pivotal to these logistical efforts. Vaccines addressing these challenges should substantially blunt the disproportionate impact COVID-19 is having on the health and economic vitality of under-resourced communities in more advanced economies and in nations with fewer financial resources.

Here, we report the development of two vaccine candidates, named AAVCOVID19-1 (AC1) and AAVCOVID19-3 (AC3), based on a unique AAV vaccine platform. We hypothesized that this methodology can address several limitations of first generation COVID vaccines. Indeed, both AAVCOVID vaccine candidates were shown in mice and NHPs to elicit robust neutralizing antibody responses following a single dose administration. This contrasts with front-runner vaccines, which require 2 injections spaced by 3 or more weeks (Folegatti et al., 2020b; Jackson et al., 2020; Logunov et al., 2020; Walsh et al., 2020). Compared to a single dose vaccine, a multiple dose regimen complicates a vaccination campaign by increasing cost, reducing compliance, and multiplying the manufacturing needs. Similarly, durable vaccine responses prolong or prevent the need for boost injections over time, resulting in similar benefits just described for single prime injection vaccines. AAVCOVID candidates have been shown in NHPs to retain peak immunogenicity for at least 5 months following a single dose injection.

The durability of other vaccine candidates remains to be determined, although a recent report on the Moderna follow up from a Phase 1 study reported the maintenance of a robust, albeit modestly declining, antibody response in 34 participants across age groups (Widge et al., 2020).

While it remains difficult to project the efficacy of the AAVCOVID vaccines based on immunological readouts alone, current data suggest an important role of neutralizing antibody responses in the prevention or mitigation of disease caused by SARS-CoV-2 (Chandrashekar et al., 2020; McMahan et al., 2020; Mercado et al., 2020; Yu et al., 2020). AAVCOVID in NHP leads to neutralizing antibody levels between 1:1,024 and 1:12,800, exceeding titers in convalescent, symptomatic, non-hospitalized patients and within the range of hospitalized and ICU convalescent COVID-19 patients. Previously reported NHP SARS-CoV-2 challenge studies indicate that neutralizing titers of greater than 1:100 are protective against disease (McMahan et al., 2020; Mercado et al., 2020; Yu et al., 2020). Further studies of the AAVCOVID vaccines are in progress that explore significant dose reductions from what is reported in this study as well as SARS-CoV-2 challenge studies that will establish whether the immune responses achieved at those lower doses laed to protective immunity.

Based on clinical observations and laboratory findings, local and systemic safety and reactogenicity was minimal in the mouse and NHP studies at all the doses tested, indicative of a favorable safety and reactogenicity profile of AAVCOVID candidates. If these findings bear out clinically, they may indicate a more limited reactogenicity compared to those seen in mRNA and adenoviral COVID-19 vaccine candidates (Corbett et al., 2020; Jackson et al., 2020; Wu et al., 2020). However, AAVrh32.33 has never been used in humans, and thus warrants additional scrutiny, particularly given the potential for use as a preventative vaccine in a healthy population where the risk-benefit ratio may substantially differ from that present in gene therapy trials. Safety concerns may be allayed given the extensive clinical experience with AAV in gene therapy applications via local - including intramuscular - or systemic routes of administration, the latter often at doses exceeding the doses proposed here by 100 to 1000-fold. These systemic studies have highlighted the potential for moderate to severe dose-related events of hepatotoxicity (Chand et al., 2020). Unlike most of the vectors pursued in gene therapy studies, AAVrh32.33 is minimally hepatotropic, particularly with the low dose intramuscular injections presented here (Figure S8D). Another concern in the use of AAV vectors as vaccines is the potential for persistent antigen expression, which is the attribute that has made them so attractive as gene therapy vectors.

Persistent antigen could lead to desensitization of the host toward the antigen through mechanisms of anergy or tolerance. Previously, in a heterologous prime-boost HIV vaccine study in NHPs, rh32.33 functionally recalled both B and T-cell responses to a greater extent than an analogous AAV8 vaccine (Lin et al., 2009). The data supports the hypothesis that AAVrh32.33 has a vaccine appropriate phenotype, unlike other AAVs, since upon IM injection, the AAV capsid acts as an adjuvant to establish a pro-inflammatory local environment that leads to extinction of transgene expression (Figure S8). The data presented in this study indicates that the biodistribution data of AAVCOVID make systemic liver toxicity following an IM injection very unlikely. Furthermore, the transgene expression data provide reassuring evidence that persistent antigen expression will also not be a limiting feature of AAVCOVID use. Taken together, these findings substantially de-risk the platform and ongoing toxicology and safety experiments are intended to further address these important issues.

An important attribute of any vaccine is its efficacy across all possible treatment populations enabling protection of the most vulnerable and thereby maximizing the potential to achieve population immunity. Here, we build on the minimal pre-existing immunity to AAVrh32.33, which should preclude significant numbers of individuals from failing to respond, as has been demonstrated for AAVs and other viral vectors for which pre-existing immunity rates are much higher (Fausther-Bovendo and Kobinger, 2014; Lin et al., 2009). Moreover, AAVCOVID induces only low-level anti-capsid responses in NHP, potentially permitting subsequent homologous boosts, as a prior study demonstrated AAV IM re-administration to be unaffected by serum AAV neutralizing antibody titers that do not exceed 1:160 (Greig et al., 2016). In addition, capsid antibody responses appear to be serotype specific and do not cross-neutralize alternative AAV serotypes that are commonly used in gene therapy. Thus, AAVCOVID immunization is unlikely to alter eligibility for a future gene therapy treatment should that ever be relevant. Lastly, AC1 enables the induction of a potent humoral immune response in mouse models of obesity and age, which in humans are associated with increased vulnerability to COVID-19 and - generally - reduced vaccine efficacy (Cuschieri and Grech, 2020; Karlsson et al., 2016; Kim et al., 2020; Park et al., 2014). While responses were reduced in 2-year-old mice as compared to younger mice of the same strain, it remains unclear how these models predict the immunogenicity in older humans, nor have studies in these models been reported elsewhere for SARS-CoV-2 vaccine candidates. Interestingly, however, AC1 potency was qualitatively distinctly higher in these models compared to that of AC3.

Some logistical attributes of the AAVCOVID vaccines that may initially seem less important biologically may, in fact, represent some of the more important advantages of this platform. Once the AAVrh32.33 vector has been proven safe in humans, its rapidly re-engineered transgene structure combined with its adaptation to large scale manufacturing and minimal cold-chain storage requirement are features that could enable its swift deployment for use in future non-COVID epidemic infections. No novel technology beyond what exists today in the AAV gene therapy manufacturing sphere is required to produce clinical grade vaccines in millions of doses. This rapid synthesis and manufacturing deployment in concert with the thermostability of AAV particles, are attractive for vaccine campaigns launched worldwide, and may be essential for those required in nations lacking transportation and sophisticated refrigeration infrastructure.

In summary, AAVCOVID is a preventative vaccine candidate for SARS-CoV-2 capable of inducing robust immunogenicity in both rodents and primates following a single injection. The safety profile in the animals studied and the level of immunogenicity compare favorably with the findings reported in studies of the COVID vaccines that have recently reported human efficacy in clinical trials. AAVCOVID vaccines are amenable to large scale production, utilizing the extensive know-how and capacity of current commercial AAV manufacturers. Biophysical properties of the vaccines indicate that a more facile distribution program can be used with AAVCOVID given the minimal cold-chain infrastructure needed.

Altogether, these data suggest that AAVCOVID is a promising preventative vaccine for SARS-CoV-2 that warrants progression to testing in human populations.

## Acknowledgements

These studies would not have been possible without the responsiveness and help of dozens of individuals within Mass Eye and Ear, Mass General, Mass General Brigham Innovation, the Gene Therapy Program at the University of Pennsylvania, Novartis Gene Therapies, Novartis Institutes for Biomedical Research, the Penn Center for Innovation, ReGenX Bio, 5AM Ventures, Aldevron, Catalent, AskBio/Viralgen, PPD, BioReliance. We thank the group of Scott Hensley, PhD for the production of fluorophore-conjugated rRBD protein used in flow cytometry studies, and Caitlyn Webb from the Grousbeck Center Gene Transfer Vector Core at Mass Eye and Ear for AAV production. Funding to this project was provided by donations from Giving/Grousbeck (Emilia Fazzalari and Wyc Grousbeck) and multiple other donors (Nathalie, Alexandre and Charles de Gunzburg; David Vargo; Julia and Mark Casady and the One Step Forward Education Foundation; Katrine S. Bosley; Tamra Gould and Howard Amster II Donor Advised Fund of the Jewish Federation of Cleveland; The Tej Kohli Foundation; Michel Plantevin; Susan Stoddart and Chris Snook; Delori Family; Annette and Dan Nova; Jennifer and Jonathan Uhrig; Lyle Howland and Jack Manning; Michelle and Bob Atchinson; Elizabeth and Phill Gross; William and Carolyn Aliski) through the Mass Eye and Ear donor network (L.H.V.); an International Fellowship from the Fundacion Alfonso Martin Escudero (N.Z.); grants from the Massachusetts Consortium for Pathogen Readiness and Mark and Lisa Schwartz (L.H.V. and A.G.S.); George Mason University Fast Grants; the Bill and Melinda Gates Foundation (L.H.V.), NIH R01 AI146779 (A.G.S.) and training grants (NIGMS T32 GM007753 for B.M.H. and NIH T32 AI007245 for J.F.); Sponsored Research Agreements from Albamunity (L.H.V and J.M.W); the U.S. Centers for Disease Control and Prevention CK000490 (E.T.R., R.C.L., R.C.C.) and an in-kind donation of AAV manufacturing services and product by Novartis Gene Therapies. The following reagent was obtained through BEI Resources, NIAID, NIH: VERO C1008 (E6), Kidney (African green monkey), Working Cell Bank, NR-596. The SARS-CoV-2 starting material was provided by the World Reference Center for Emerging Viruses and Arboviruses (WRCEVA), with Natalie Thornburg as the CDC Principal Investigator. Avicel RC-591 was kindly provided by DuPont Nutrition & Health.

## Author contributions

Conceptualization, N.Z., W.D., R.H., M.R.B., J.M.W., and L.H.V.; Methodology, N.Z., W.D., U.B., J.A.C., C.D.D., S.Y., R.H. and L.H.V.; Validation, N.Z., W.D., U.B., W.Q. and E.H.; Formal Analysis, N.Z., J.A.C., J.S., M.B.P., J.J.K and L.H.V.; EZ provided the phylogenetic analysis in Fig1B; Investigation, N.Z., W.D., U.B., J.A.C., J.S., R.E., C.D., D.M., W.Q., E.H., A.C., C.D.D., M.B.P., J.J.K., D.L., M.K., A.S., N.S. and R.J.; Resources, A.C., R.C.L., R.C.C., A.S., J.F., B.M.H., D.T.K, R.C., E.T.R., A.G.S., J.M.W. and L.H.V.; Writing – Original Draft, N.Z., M.W.F. and L.H.V.; Writing – Review and Editing, N.Z., W.D., U.B., J.A.C., J.S., D.M., M.B.P., J.J.K., A.R., R.C., M.M., E.T.R., B.P., J.M.W. and L.H.V.; Visualization, N.Z., U.B., M.B.P. and J.J.K.; Supervision, N.Z., K.T.M., C.D.D., D.T.K., A.H., A.G., S.Y., R.H., B.P., M.R.B., M.W.F., J.M.W., and L.H.V.; Project Administration, N.Z., K.T.M., A.R., D.T.K., R.C., M.M., A.H. and B.P.; Funding Acquisition: M.W.F, J.M.W. and L.H.V.

## Declaration of Interests

JMW is a paid advisor to and holds equity in Scout Bio and Passage Bio; he holds equity in Surmount Bio; he also has sponsored research agreements with Amicus Therapeutics, Biogen, Elaaj Bio, Janssen, Moderna, Passage Bio, Regeneron, Scout Bio, Surmount Bio, and Ultragenyx, which are licensees of Penn technology. LHV and JMW are inventors on patents that have been licensed to various biopharmaceutical companies and for which they may receive payments. MWF is a paid consultant to 5AM Ventures and to Mitobridge/Astellas. LHV is a paid advisor to Novartis, Akouos, Affinia Therapeutics and serves on the Board of Directors of Affinia, Addgene and Odylia Therapeutics. LHV holds equity in Akouos and Affinia and receives sponsored research funding from Albamunity Inc. to which he is an unpaid consultant.

## MATERIAL AND METHODS

### Vaccine candidates

Two AAV-based vaccine candidates were tested: AAVCOVID19-1 (AC1) and AAVCOVID19-3 (AC3) (Figure 1A) (plasmid sequences submitted to GenBank, submission ID #2410698). AC1 is an AAVrh32.33 vector that expresses the codon optimized, pre-fusion stabilized (furin cleavage site mutated to G682SAS685 and P986P987 substitutions) full length SARS-CoV-2 Spike protein under the control of an SV40 promoter. AC1 carries a short SV40 polyadenylation signal (poly-A). AC3 is an AAVrh32.33 that carries the secreted S1 subunit of SARS-CoV-2 Spike with the tissue plasminogen activator signal peptide (tPA-SP) whose expression is driven by the CMV promoter. AC3 has two more regulatory elements: a woodchuck hepatitis virus posttranscriptional regulatory element (WPRE) and the bovine growth hormone polyadenylation signal (poly-A).

### Small-scale production of vaccine candidates

Research-grade, high-titer vectors were produced, purified, and titrated by the MEEI/ SERI Gene Transfer Vector Core (https://www.vdb-lab.org/vector-core/). Small-scale vector preparations were generated by polyethylenimine or PEI (Polysciences, Cat #24765-2) triple transfection of AC1 or AC3 ITR-flanked transgene, pKan2/rh32.33 (AAV2 rep and AAVrh32.33 capsid construct), and pALD-X80 adenoviral helper plasmid in a 1:1:2 ratio, respectively, in HEK293 cells. DNA was transfected in 10-layer HYPERFlasks using a PEI-Max/DNA ratio of 1.375:1 (v/w). 3 days after transfection, vectors were harvested from the HYPERFlasks using Benzonase (EMD Millipore, catalog no. 1016970010) to degrade DNA/RNA. 24 hours after harvesting, the vectors were concentrated by tangential flow filtration and purified by iodixanol gradient ultracentrifugation as previously described (Lock et al., 2010). Vaccine candidates were quantified by ddPCR according to a previously published protocol (Sanmiguel et al., 2019). Capsid stability was assessed by AAV-ID (Pacouret et al., 2017).

### Large-scale manufacturing of vaccine candidates

AAVCOVID candidates were produced at larger scale via standard AAV production processes by Novartis Gene Therapies, following their stablished protocol with only minimal modifications to adjust to the AAVrh32.33 technology. Briefly, AC1 and AC3 were produced via three plasmid transfection (AC1 or AC3 ITR-flanked transgene, pKan2/rh32.33 (AAV2 rep and AAVrh32.33 capsid construct), and pALD-X80 adenoviral helper plasmid) in an iCellis500 bioreactor (Pall Biosciences). Following cell lysis and lysate clarification, tangential flow filtration (TFF) was conducted to achieve volume reduction. The TFF retentate was next enriched for AAV particles on a cation exchange chromatography column (BIA Separations, Sartorius). The eluate was concentrated, and buffer exchanged through an additional TFF step, before CsCl ultracentrifugation to separate genome containing versus empty AAV particles. Finally, formulation (buffer: 20 mM tris (pH 8.1 ± 0.1), 1 mM magnesium chloride (MgCl2), 200 mM sodium chloride (NaCl) and 0.005% poloxamer 188) was achieved through TFF before bulk drug substance was filtered.

### TaqMan™ Assay Design for SARS-CoV-2 Spike detection

The codon optimized SARS-CoV-2 receptor binding domain (RBD) of AAVCOVID vaccine candidates was used as a target for droplet digital PCR (ddPCR)/real-time PCR (qPCR) quantifications. The sequence was checked for secondary structures using the mfold application of the UNAfold software package (Zuker, 2003) at the PCR annealing temperature and TaqMan buffer salt concentrations.Internal repeats were avoided by maping against the entire codon optimized SARS-CoV-2 S gene of AAVCOVID candidates using the REPuter application (Kurtz et al., 2001). The 5’-end of the gene was selected as PCR target based on these analyses. The oligo sequences used were the following: forward primer, GTGCAGCCAACCGAG (0.43μM final concentration); reverse primer, ACACCTCGCCAAATGG (1.125μM final concentration), and TaqMan® probe 6FAM-TCTATCGTGCGCTTTC-MGBNFQ (0.25μM final concentration). The final concentration and Tm’s of primers were determined using the DINAMelt application of the UNAfold software package (Markham and Zuker, 2005, 2008) and set to hybridize the target with a Tm of just under 60°C (59.0-59.9°C) for high specificity. The PrimerExpress™ software (Applied Biosystems™) was used to determine the Tm of the MGB probe (Kutyavin et al., 2000). The resulting 67 bp amplicon was inspected for specificity via NCBI BLAST® using the somewhat similar algorithm in the suite against human, NHP, mouse, ferret, and betacoronavirus databases and determined to be highly specific for our vaccine candidates. No significant matches were found against the RBD oligonucleotides used.

### In vitro expression studies

10^5^ HEK293 cell/well were seeded in 12-well plates (Corning, MA, USA) plates and incubated at 37°C overnight. The following day, cells were transfected with 2 µg of AAVCOVID19-1 (pAC1) and AAVCOVID19-3 (pAC3) plasmids using PEI-Max. Cells were harvested 24 and 72 hours after transfection for mRNA and western blot (WB) expression analyses, respectively. In addition, 5×10^4^ HuH7 cell/well were seeded in 12-well plates and incubated overnight at 37°C. On the following day, adenovirus 5 WT (Ad5) was added to the cells at a MOI of 20 pfu/cell. 2 hours later, media was removed and cells infected with a MOI of 5×10^5^ of AC1 or AC3. Cells were harvested 72 hours later for WB analysis.

Transfection and transduction samples were also collected for RNA gene expression analyses. Total RNA was extracted via Trizol™ reagent (Invitrogen™) and quantified using a Qubit™ fluorometer (Invitrogen™). 7.5µg of Total RNA was DNase-I treated using the Turbo DNA-*free*™ kit (Invitrogen™). About 1.4µg of DNase-treated total RNA was set aside for reverse transcription against (-)RT controls using the high capacity cDNA reverse transcription kit (Thermo Fisher™). Codon optimized RBD gene expression was assessed against a cells only control using qPCR and normalized to human 18S rRNA gene levels by the delta delta Ct method (Livak and Schmittgen, 2001).

### Detection of Spike antigens by Western Blot (WB)

Cell lysates were obtained by diluting cell pellets in NuPAGE™ LDS Sample Buffer (4X) (Thermo Fisher Scientific, Cat# NP0007) and incubating at 99°C for 5 minutes,, separated by electrophoresis in NuPAGE 4-12% polyacrylamide gels (Thermo Fisher Scientific, Cat#NP0321PK2) and then transferred to PVDF membranes. The membranes were probed with an anti-SARS-CoV-2 RBD rabbit polyclonal antibody (Sino Biological Inc., 40592-T62) followed by a goat anti-rabbit HRP-conjugated secondary antibody (Thermo Fisher Scientific, Cat# A16110, RRID AB_2534782). Membranes were developed by chemiluminescence using the Immobilon Western Chemiluminescent HRP Substrate (Millipore, Cat# WBKLS0500) and recorded using ChemiDoc MP Imaging System (Bio-Rad). An anti-GAPDH antibody (Cell Signaling Technology Cat# 2118, RRID:AB_561053) was used as loading control.

### Mouse studies

All the mouse studies were performed in compliance with the Schepens Eye Research Institute IACUC. BALB/c, C57BL/6 or C57BL/6 diet-induced obese (DIO) animals were intramuscularly (right gastrocnemius muscle) treated at 10^10^ gc/mouse or 10^11^ gc/mouse. Animals were kept in standard diet and C57BL/6 DIO were fed a high-fat diet (Research Diets, Cat#D12492i). Serum samples were obtained by submandibular bleeds for humoral immune response analyses. At necropsy, several tissues were collected for analysis of vector presence and transgene expression.

### NHP study

All animal procedures were approved by the Institutional Animal Care and Use Committee of the Children’s Hospital of Philadelphia. Rhesus macaques (Macaca mulatta) that screened negative for viral pathogens including SIV (simian immunodeficiency virus), STLV (simian-T-lymphotrophic virus), SRV (simian retrovirus), and B virus (macacine herpesvirus 1) were enrolled on the study. Animals were housed in an AAALAC International-accredited nonhuman primate research in stainless-steel squeeze back cages, on a 12-hour timed light/dark cycle, at temperatures ranging from 64-79°F (18-26°C). Animals received varied enrichment such as food treats, visual and auditory stimuli, manipulatives, and social interactions throughout the study. Four 3 to 7 year-old Rhesus macaques (*Macaca mulatta*) were treated with the clinical candidates (n=2/vector, 1 female and 1 male) intramuscularly at a dose of 10^12^ gc/animal. Serum and PBMC samples were obtained in regular intervals for several analyses of immunogenicity against SARS-CoV-2 Spike and AAVrh32.33. Serum chemistry, hematology, and coagulation analyses were performed by Antech Diagnostics. Serum was also collected for cytokine analyses which were performed by the University of Pennsylvania’s Human Immunology Core using a Non-Human Primate Cytokine Panel kit (MilliporeSigma, Cat# PCYTMG-40K-PX23) on a Bio-Plex 200 instrument (Bio-Rad) according to the manufacturer’s protocol.

### Human samples

Blood was collected from 60 patients with nasopharyngeal PCR-confirmed SARS-CoV-2 infection stratified by disease severity. Plasma was separated and stored at negative 80°C until assessed. Human subject investigation was approved by the institutional Review Board of the Massachusetts General Hospital.

### SARS-CoV-2 Spike-binding antibody detection ELISA

Nunc MaxiSorp^TM^ high protein-binding capacity 96 well plates (Thermo Fisher Scientific, Cat# 44-2404-21) were coated overnight at 4 °C with 1µg/ml SARS-CoV-2 RBD, SARS-CoV-2 ectodomain (LakePharma, Cat# 46328) or SARS-CoV-1 RBD diluted in phosphate-buffered saline (PBS). The next day the plates were washed with PBS-Tween 20 0.05% (Sigma, Cat# P2287-100ML) using the Biotek 405 TS Microplate washer. Each plate was washed five times with 200 µl wash buffer and then dried before the next step. Following the first wash, 200 µl of Blocker Casein in PBS (Thermo Fisher Scientific, Cat# 37528) were added to each well and incubated for 2 hours at RT. After blocking, serum samples were serially diluted in blocking solution starting into 1:100 dilution. After an hour of incubation, the plates were washed and 100 µl of secondary Peroxidase AffiniPure Rabbit Anti-Mouse IgG (Jackson ImmunoResearch, Cat# 315-035-045, RRID: AB_2340066) antibody diluted 1:1000 in blocking solution or rabbit Anti-Monkey IgG (whole molecule)-Peroxidase antibody (Sigma-Aldrich Cat# A2054, RRID:AB_257967) were added to each well. After one hour of incubation at room temperature, the plates were washed and developed for 3.5 min with 100 µl of Seracare SureBlue Reserve^TM^ TMB Microwell Peroxidase Substrate solution (SeraCare, Cat# 53-00-03). The reaction was then stopped with 100 µl Seracare KPL TMB Stop Solution (SeraCare, Cat# 50-85-06). Optical density (OD) at 450 nm was measured using a Biotek Synergy H1 plate reader. The titer was the reciprocal of the highest dilution with absorbance values higher than four times the average of the negative control wells. For mouse serum SARS-CoV-2 RBD-specific antibody isotyping, the same ELISA was performed but using the secondary antibodies from SBA Clonotyping System-HRP kit (SouthernBiotech, 5300-05, RRID:AB_2796080) diluted accordingly to manufacturer’s instructions.

For NHP isotyping ELISA, plates were coated overnight at 4°C with 20ng/well of SARS-CoV-2 Spike protein ectodomain (LakePharma, Cat# 46328) as the capture antigen. Then, the wells were blocked with 5% milk in PBS-Tween 20 0.05% for 2h and incubated for additional 2h with 50 μl serially diluted serum samples. Then, horseradish peroxidase (HRP)-conjugated secondary antibody against rhesus IgG1 (NIH Nonhuman Primate Reagent Resource supported by AI126683 and OD 010976 Cat# PR-7110, RRID:AB_2819310) and IgG4 (NIH Nonhuman Primate Reagent Resource supported by AI126683 and OD 010976 Cat# PR-7180, RRID:AB_2819322) for 1 h. After every incubation step, the plates were washed three times with PBS-Tween 20 0.05%. After color development, OD at 450nm was determined using Biotek Synergy H1 plate reader. The titer was the reciprocal of the highest dilution with absorbance values higher than four times the average of the negative control wells.

### Lenti-SARS2 pseudovirus production and titration

Lenti-SARS2 was produced based on a published protocol (Crawford et al., 2020). Specifically, 50% confluent HEK293T cells were seeded 24 hours prior to transfection in 15 cm plates. The next day, 18µg of psPAX2, 9µg of pCMV-SARS2-RRAR_ILR_gp41 and 29µg of pCMV-Lenti-Luc plasmids were mixed in 3.6mL of Opti-MEM™ I Reduced Serum media (Gibco, Cat# 31985070) along with 144 µL of PEI Max 40K (1 mg/mL, pH: 6.9-7.1) and mixed thoroughly. The mixture was incubated for 20 minutes at room temperature. Media on cells was aspirated and serum-free DMEM was added to the cells. After 20 mins, the DNA-PEI mixture was added dropwise to the plate and incubated overnight at 37°C with 5% CO2. The next day, media was replaced with DMEM with 10% FBS. After 48 hours, the media was collected in a 50 mL conical and centrifuged at 4,000 rpm at 4°C for 5 minutes to remove cell debris. The supernatant was collected and filtered through 0.45 µm filter, aliquoted and stored at −80°C.

For titration of the pseudovirus, HEK293T cells expressing ACE2 were seeded at 1.5×10^4^ cells/well in poly-L-Lysine (0.01%) coated 96-well black plates (Thermo Fisher Scientific, Cat# 3904) one day before titration. On the next day, the media was changed to 50 µL DMEM+10%FBS containing filtered Hexadimethrine bromide at a final concentration of 10 µg/mL. 2-fold serial dilutions (up-to 15 dilutions) of the viral stocks (50µL) were added to the plate in 6 replicates each and incubated for 48 hours. After 48 hours, cells were lysed with Reporter Lysis Buffer (Promega, Cat# E4030). These plates were frozen at − 80°C for 60 minutes. Thereafter, they were thawed at 37°C for 20 mins before starting the luciferase readout. For luciferase substrate buffer the following reagents were mixed; Tris-HCl buffer at 0.5 M, ATP at 0.3 mM, MgCl2 at 10 mM, Pierce™ Firefly Signal Enhancer (Thermo Fisher Scientific, Cat#16180), D-luciferin 150µg/mL (PerkinElmer, Cat# 122799). Biotek Synergy H1 Plate reader was used for luminescence readout. For pseudovirus neutralization assay, a final dilution of the virus stock targeting relative luminescence units (RLU) of 1800-1100 was used which was approximately 200-fold higher than background signal obtained in untreated cells.

### Pseudovirus Neutralization Assay

HEK293T cells expressing ACE2 were seeded at 1.5×10^4^ cells/well in poly-L-Lysine (0.01%) coated 96-well black plates. The following day, 50µL of DMEM+10% FBS media containing Hexadimethrine bromide (final concentration 10 µg/mL) was added to the cells. Serum samples were heat-inactivated at 56°C for 1 hour. Serum samples were then serially diluted (2-fold) for 10 dilutions in DMEM with 10% FBS with initial dilution of 1:40 for mouse serum and 1:10 dilution for NHP serum. Thereafter, Lenti-SARS2 pseudovirus was added to each dilution and incubated at 37°C for 45 minuntes. The serum and virus mixture was added to the cells and incubated at 37°C with 5% CO2 for 48 hours. An anti-SARS-CoV-2 Spike monoclonal neutralizing antibody (GenScript, Cat# 6D11F2) was used as a positive control. Cells without serum and virus were used as negative control. After 48 hours, cells were lysed and luciferase measured as described above. Neutralizing antibody titers or 50% inhibitory concentration in the serum sample (EC50 or ID50) were calculated as the reciprocal of the highest dilution showing less RLU signal than half of the average RLU (maximum infectivity) of Virus Control group (cells + virus, without serum).

### Plaque Reduction Neutralization Test (PRNT) of SARS-CoV-2

Depending on the volume available, mouse or NHP sera were serially diluted two-fold from an initial dilution of either 1:12.5 or 1:25 for ten dilutions in Dulbecco’s Phosphate Buffered Saline (DPBS, Gibco). Each dilution was incubated at 37°C and 5% CO2 for 1 hour with an equal volume of 1000 plaque forming units/ml (PFU/ml) of SARS-CoV-2 (isolate USA-WA1/2020) diluted in DMEM (Gibco) containing 2% fetal bovine serum (Gibco) and antibiotic-antimycotic (Gibco). Controls included DMEM containing 2% fetal bovine serum (Gibco) and antibiotic-antimycotic (Gibco) only as a negative control, 1000 PFU/ml SARS-CoV-2 incubated with DPBS, and 1000 PFU/ml SARS-CoV-2 incubated with DMEM. Two hundred microliters of each dilution or control were added to confluent monolayers of NR-596 Vero E6 cells in triplicate and incubated for 1 hour at 37°C and 5% CO2. The plates were gently rocked every 5-10 minutes to prevent monolayer drying. The monolayers were then overlaid with a 1:1 mixture of 2.5% Avicel® RC-591 microcrystalline cellulose and carboxymethylcellulose sodium (DuPont Nutrition & Biosciences) and 2X Modified Eagle Medium (Temin’s modification, Gibco) supplemented with 2X antibiotic-antimycotic (Gibco), 2X GlutaMAX (Gibco) and 10% fetal bovine serum (Gibco).

Plates were incubated at 37°C and 5% CO2 for 2 days. The monolayers were fixed with 10% neutral buffered formalin and stained with 0.2% aqueous Gentian Violet (RICCA Chemicals) in 10% neutral buffered formalin for 30 minutes, followed by rinsing and plaque counting. The half maximal inhibitory concentrations (EC50 or ID50) were calculated using GraphPad Prism 8.

### IFN-γ and IL-4 ELISPOT assay and cytokine secretion in murine splenocytes

Splenocytes were obtained by grinding murine spleens with 100 μm cell strainers, followed by treatment with Ammonium Chloride-Potassium (ACK) lysis buffer (Gibco) to lyse the red blood cells. The isolated cells were then suspended in complete RPMI-1640 medium (Gibco) supplemented with 10% FBS and counted for the following experiments.

IFN-γ and IL-4 ELISPOT for mice was measured as previously described (Wang et al., 2019). Briefly, 96-well PVDF plates (Millipore) were pre-coated with 10 μg/ml anti-mouse IFN-γ ELISPOT capture antibody (BD Biosciences Cat# 551881, RRID:AB_2868948) or 4 μg/ml anti-mouse IL-4 ELISPOT capture antibody (BD Biosciences Cat# 551878, RRID:AB_2336921) at 4°C overnight, and then blocked with complete RPMI-1640 medium for 3 hours at 37°C. One million of freshly isolated splenocytes were seeded into the precoated plates and stimulated with S1 and S2 peptides pools (GenScript) with a final concentration of 1 μg/ml of each peptide diluted in RPMI-1640 supplemented with 10% FBS and incubated for 48 hours at 37°C with 5% CO2. Each peptide pool, consisting of 15-mers peptides overlapping by 10 amino acids, spanning the entire SARS-CoV-2 Spike protein S1 or S2 subunits.

Control wells contained 5×10^5^ cell stimulated with DMSO diluted in RPMI-1640 supplemented with 10% FBS (negative control) or 2 μg/ml concanavalin A (positive control). Subsequently, the plates were washed and incubated with biotin-conjugated mouse IFN-γ ELISPOT Detection Antibody (BD Biosciences Cat# 551881, RRID:AB_2868948) and 4 μg/ml biotin-conjugated mouse IL4 detection antibody (BD Biosciences Cat# 551878, RRID:AB_2336921) at room temperature for 3 hours and followed by streptavidin-HRP (dilution 1:1000, Sigma-Aldrich, Cat# 18-152) for 45 minutes. After washing, 100 μL/well of NBT/BCIP substrate solution (Promega, Cat# S3771) were added and developed for 15–30 min until distinct spots emerged. The cytokine-secreting cell spots were imaged and counted on AID EliSpot reader (Autoimmun Diagnostika GmbH).

2×10^6^ freshly isolated splenocytes were seeded into 96-well plates and stimulated with 1 µg/ml of peptides from S1 and S2 pool as described previously at 37 °C for 48 hours. Then the supernatants were collected and cytokine levels were measured by a Luminex cytokine assay by SBH Sciences.

### Detection of circulating S1 in NHP sera by Sandwich ELISA

A monoclonal anti-SARS-CoV-2 RBD capture antibody (GenScript, Cat# 5B7D7) was coated on Nunc Maxisorp ELISA plates (Thermo Fisher Scientific, Cat# 44-2404-21) at 2.5 µg/mL final concentration in Sodium Bicarbonate buffer (Sigma-Aldrich, Cat# SRE0034). The plate was incubated at 4°C overnight. All washes were performed 5X with PBS-Tween-20 0.05%. On the following day, plates were washed and blocked for 2 hours with Casein Buffer (Thermo Fisher Scientific, Cat# 37528). Then, NHP sera were added in duplicates at 1:5 dilution in blocking buffer. A blank consisting of the blocking buffer and a standard curve ranging from 5000 pg/mL to 78.25 pg/mL of S1 antigen (GenScript, Cat# Z03501) in blocking buffer were also added in duplicates on the plate followed by incubation at room temperature for 1 hour. Then, biotinylated detection antibody (GenScript, Cat# 5E10G8-Biotin) was added at 1 µg/mL final concentration in blocking buffer and plate was incubated at room temperature for 1 hour. Finally, 1:5000 final dilution of Streptavidin-HRP (Sigma-Aldrich, Cat# 18-152) was added to the plate. After completing incubation of 1 hour at room temperature, the plate was washed. 100 µL of TMB substrate (SeraCare, Cat# 5120-0081) was added to plates and color was developed for 3 mins 30 secs, after which 100 µL of Stop Solution (SeraCare, Cat# 5150-0021) was added to stop the reaction and plates were read at 450 nm and 670 nm using Biotek Synergy H1 hybrid plate reader. Absorbance at 670 nm was subtracted from 450 nm, and then corrected with absorbance of the blank. Linear regression was used to calculate the standard curve formula and S1 concentration (pg/mL) was calculated by extrapolation.

### IFN-γ ELISPOT Assay in NHP PBMCs

Peripheral blood T cell responses against AC1, AC3 and the AAVrh32.33 capsid were measured by interferon gamma (IFN-γ) enzyme-linked immunosorbent spot (ELISPOT) assays according to previously published methods (Calcedo et al., 2018). Peptide libraries specific for AAVrh32.33 capsid as well as the AC1 and AC3 transgenes were generated (15-mers with a 10 amino acid overlap with the preceding peptide; Mimotopes, Australia). More specifically, the AAVrh32.33 capsid peptide library was divided into three peptide pools, A, B and C. Pool A contained peptides 1-50, Pool B peptides 51-100 and Pool C peptides 101-145. For the AC1 and AC3 peptide libraries, peptides specific to each protein were pooled separately from those peptide sequences shared between the two proteins. The AC1 peptide library contained Pool A (peptides 1-2, 136-173); Pool B (peptides 174-213); and Pool C (peptides 214-253).

The AC3 Peptide Library consisted of Pool A only (peptides 254-257). The AC1 & AC3 Shared Peptides also contained three peptide pools; Pool A (peptides 258-259; 3-44), Pool B (peptides 45-90) and Pool C (peptides 91-135). Peptides were dissolved in dimethyl sulfoxide (DMSO) at a concentration of 100 mg/mL, pooled, aliquoted and stored at −80°C. They were used at a final concentration in the assay of approximately 2 µg/mL. The positive response criteria for the IFN-γ ELISPOT was greater than 55 spot forming units (SFU) per million cells and at least three times greater than the negative control values.

### PBMC stimulation, flow cytometry and gating strategy

Cryopreserved peripheral blood mononuclear cells (PBMC) were thawed and rested overnight in sterile R10 media (RPMI 1640, Corning), supplemented with 10% fetal bovine serum (Gemini Bio-Products), Penicillin/Streptomycin and L-Glutamine; plus 10 U/mL DNAse I (Roche Life Sciences) at 37°C, 5% CO2 and 95% humidity incubation conditions. PBMC were stimulated at 200 ul final volume in sterile R10 media. Peptide concentrations for stimulation conditions were 2 µg/ml for AC1/AC3 shared peptide pool A, B and C and AAVrh32.22 peptide pool A, B and C. Co-stimulation was added with peptides: 1μg/mL anti-CD49d (Clone 9F10, BioLegend Cat# 304301, RRID:AB_314427) and CD28-ECD (Clone CD28.2, Beckman Coulter Cat# 6607111, RRID:AB_1575955) at the start of stimulation. Positive control samples were stimulated using Staphylococcal Enterotoxin B (SEB, List Biological Laboratories) at 1μg/mL. CD107a BV650 (clone H4A3, BioLegend Cat# 328643, RRID:AB_2565967) was added at the start of stimulation. Brefeldin A (1μg /mL) (Sigma-Aldrich) and monensin (0.66μL/mL) (BD Biosciences) were added one hour after initiation of stimulation. Cells were incubated under stimulation conditions for a total of 9 hours.

All following incubations were performed at room temperature. Cells were stained for viability exclusion using Live/Dead Fixable Aqua for 10 minutes, followed by a 20-minute incubation with a panel of directly conjugated monoclonal antibodies diluted in equal parts of fluorescence-activated cell sorting (FACS) buffer (PBS containing 0.1% sodium azide and 1% bovine serum albumin) and Brilliant stain buffer (BD Biosciences). Fluorophore-conjugated recombinant RBD protein produced by the Hensley Lab (University of Pennsylvania) was used to identify RBD-binding B cells during the surface antibody stain. Cells were washed in FACS buffer and fixed/permeabilized using the FoxP3 Transcription Factor Buffer Kit (eBioscience), following manufacturer’s instructions. Intracellular staining was performed by adding the antibody cocktail prepared in 1X permwash buffer for 1 hour. Stained cells were washed and fixed in PBS containing 1% paraformaldehyde (Sigma-Aldrich) and stored at 4°C in the dark until acquisition. All flow cytometry data were collected on a BD LSR II or BD FACSymphony A5 cytometer (BD Biosciences). Data were analyzed using FlowJo software (versions 9.9.6 and 10.6.2, Tree Star).

The following antibodies were used: PD1 BV421 (clone EH12.2H7, BioLegend Cat# 329919, RRID:AB_10900818), CD14 BV510 (clone M5E2, BioLegend Cat# 301842, RRID:AB_2561946) and APC-Cy7 (clone M5E2, BioLegend Cat# 301819, RRID:AB_493694), CD16 BV510 (clone 3G8, BioLegend Cat# 302048, RRID:AB_2562085) and APC-Cy7 (clone 3G8, BioLegend Cat# 302017, RRID:AB_314217), CD20 BV510 (clone 2H7, BioLegend Cat# 302339, RRID:AB_2561721) and BV650 (clone 2H7, BioLegend Cat# 302335, RRID:AB_11218609), CD69 BV605 (clone FN50, BioLegend Cat# 310937, RRID:AB_2562306), CD21 PECy7 (clone Bu32, BioLegend Cat# 354911, RRID:AB_2561576), CD4 BUV661 (clone SK3, BD Biosciences Cat# 612962, RRID:AB_2870238), CD95 BUV737 (clone DX2, BD Biosciences Cat# 612790, RRID:AB_2870117), CD8 BUV563 (clone RPA-T8, BD Biosciences Cat# 612914, RRID:AB_2870199), KI67 BV786 (clone B56, BD Biosciences Cat# 563756, RRID:AB_2732007), IL2 PE (clone MQ1-17H12, BD Biosciences Cat# 554566, RRID:AB_395483), IFNγ BV750 (clone B27, BD Biosciences Cat# 566357, RRID:AB_2739707), CD3 BUV805 (clone SP34-2, BD Biosciences Cat# 742053, RRID:AB_2871342), Granzyme B AF700 (clone GB11, BD Biosciences Cat# 560213, RRID:AB_1645453), CD3 APC-Cy7 (clone SP34-2, BD Biosciences Cat# 557757, RRID:AB_396863), IgM PECy5 (clone G20-127, BD Biosciences Cat# 551079, RRID:AB_394036), CD27 BV421 (clone M-T271, BD Biosciences Cat# 562513, RRID:AB_11153497), HLA-DR BV605 (clone G46-6, BD Biosciences Cat# 562844, RRID:AB_2744478), CD80 BV786 (clone L307.4, BD Biosciences Cat# 564159, RRID:AB_2738631), CXCR3 AF488 (clone 1C6, BD Biosciences Cat# 558047, RRID:AB_397008), CXCR5 SB702 (clone MU5BEE, Thermo Fisher Scientific Cat# 67-9185-42, RRID:AB_2717183), Tbet PerCP-Cy5.5 (clone 4B10, Thermo Fisher Scientific Cat# 45-5825-82, RRID:AB_953657), CD11c PECy5.5 (clone 3.9, Thermo Fisher Scientific Cat# 35-0116-42, RRID:AB_11218511), TNFα PE-Cy7 (clone Mab11, Thermo Fisher Scientific Cat# 25-7349-41, RRID:AB_1257208), and polyclonal anti-IgD PE Tx Red (SouthernBiotech Cat# 2030-09, RRID:AB_2795630).

First, to ensure that only live single cells were analyzed from PBMCs, forward scatter height (FSC-H)-versus-forward scatter area (FSC-A) and side scatter area (SSC-A)-versus-FSC-A plots were used to exclude doublets and focus on singlet small lymphocytes. Dead cells were excluded by gating on cells negative for the viability marker Aqua Blue. For T cell function analysis, monocytes, B cells and NK cells were excluded via the CD14/19/16 dump gate. CD4^+^ and CD8^+^ T lymphocytes were gated within CD3^+^ cells. To determine the memory phenotype, CD28 versus CD95 were used, and naïve T cells were excluded from the analysis.

For B cell analysis, B cells were identified as CD20^+^ and CD3/CD14/CD16^-^. Memory B cells were defined as CD27^+^ or CD27^-^IgD^-^.

### AAV Neutralizing Antibody (NAb) Assay

NAb responses against AAV1, AAV2, AAV5, AAV8, AAV9 and AAVrh32.33 capsids were measured in serum using an in vitro HEK293 cell-based assay and LacZ expressing vectors (Vector Core Laboratory, University of Pennsylvania, Philadelphia, PA) as previously described (Calcedo et al., 2018). The NAb titer values are reported as the reciprocal of the highest serum dilution at which AAV transduction is reduced 50% compared to the negative control. The limit of detection of the assay was a 1:5 serum dilution.

### Biodistribution/Gene Expression Studies

Tissue collection was segregated for genomic DNA (gDNA) or total RNA work by QIASymphony nucleic acid extraction with the aim of filling up 96-well plates of purified material. A small cut of frozen tissue (∼ 20 mg) was used for all extractions with the exception of gDNA purifications from spleen (1-2 mg). Tissues were disrupted and homogenized in QIAGEN Buffer ATL (180 μL) and lysed overnight at 56°C in the presence of QIAGEN Proteinase K (400 μg) for gDNA, or directly in QIAGEN® Buffer RLT-Plus in the presence of 2-mercaptoethanol and a QIAGEN anti-foaming agent for total RNA purification. Tissue lysates for gDNA extraction were treated in advance with QIAGEN RNase A (400 µg), while tissue homogenates for RNA extraction were DNase-I treated *in situ* in the QIASymphony® during the procedure. Nucleic acids were quantified only if necessary, as a troubleshooting measure.

Purified gDNA samples were diluted 10-fold and in parallel into Cutsmart-buffered BamHI-HF (New England Biolabs) restriction digestions in the presence of 0.1% Pluronic F-68 (50μL final volume) that ran overnight prior to quantification. Similarly, DNase-I-treated total RNAs were diluted 10-fold into cDNA synthesis reactions (20 μL final volume) with or without reverse transcriptase using the High Capacity cDNA Reverse Transcription Kit (Thermo Fisher™). For ddPCR (gDNA or cDNA) or qPCR (cDNA), 2 μL of processed nucleic acids were used for quantification using Bio-Rad™ or Applied Biosystems™ reagents, respectively, in 20 μL reactions using default amplification parameters without an UNG incubation step. All the studies included negative control (PBS) groups for comparison. The significantly small variance of multiple technical replicates in ddPCR justified the use of a single technical replicate per sample and no less than three biological replicates per group, gender, or time point. coRBD signal for ddPCR and vector biodistribution (gDNA) was multiplexed and normalized against the mouse transferrin receptor (Tfrc) gene TaqMan™ assay using a commercial preparation validated for copy number variation analysis (Thermo Fisher Scientific). Likewise, coRBD signal for ddPCR and gene expression analysis was multiplexed and normalized against the mouse GAPDH gene, also using a commercial preparation of the reference assay (Thermo Fisher Scientific). Target and reference oligonucleotide probes are tagged with different fluorophores at the 5’-end which allows efficient signal stratification. For qPCR, coRBD and mGAPDH TaqMan assays were run separately to minimize competitive PCR multiplexing issues prior to analysis and delta delta Ct normalization (Livak and Schmittgen, 2001). The limit of detection of the assay was 10 copies/reaction, therefore, wells with less than 10 copies were considered negative.

### Phylogenetic Analysis

First, fourteen representative AAV capsid sequences were aligned by Clustal Omega (Sievers and Higgins, 2018). Substitution models and model parameters were statistically compared (120 in total) through ProtTest3 (Darriba et al., 2011; Guindon and Gascuel, 2003), and the Le, Gascuel Model (Le and Gascuel, 2008) was selected based on the Aikake Information Criterion (AIC). Additionally, amino acid frequencies were determined empirically through the alignment (+F parameter) and evolutionary rates among sites were allowed to vary within five categories by modeling variability with a discrete Gamma distribution (+G parameter), again selected through AIC. A maximum-likelihood approach was then used to infer the evolutionary relationships among the included sequences using MEGA X (Kumar et al., 2018) and the resultant phylogeny rooted along the midpoint of the branch between AAV4 and AAV5 for purposes of visualization. A sequence identity matrix was computed, and the resultant table was used to annotate the phylogeny by percent identity.

### Graphs and statistical analysis

GraphPad Prism 8 was used for graph preparation and statistical analysis. Data were represented as mean ± standard deviation (SD). Groups were compared between them by One-way ANOVA and Tukey’s tests in studies with more than two groups and n≥10, and Kruskal Wallis and Dunn’s testes were used if n<10. Two groups were compared between them using Student’s t test (if n≥10) or Mann Whitney’s U (if n<10). Pearson’s correlation coefficient was calculated to assess correlation.

## SUPPLEMENTAL FIGURE LEGENDS

**Figure S1. Related to Figure 1. Expression of AC1 and AC3 in vitro.** (A) coRBD mRNA expression relative to human 18S rRNA in HEK293 cells transfected with 1 µg of the ITR-containing pAC1 or pAC3 plasmids or transduced with 1×10^5^ or 5×10^5^ gc/cell of AC1 or AC3 24h after treatment. Ctr: untreated cells. (B) Unedited WB image shown in Figure 1G. Red rectangle indicate the part of the gel represented in Figure 1G.

**Figure S2. Related to Figure 2. Short term antibody kinetics in mice**. (A) RBD-binding IgG antibody titers measured weekly during the first month after vaccination in BALB/c (n=5 females and 5 males) treated IM with two doses of AC1 and AC3. Data are represented as mean ± SD. Groups were compared by one-way ANOVA and Tukey’s post-test. * p<0.05, ** p<0.01. (B) Seroconversion rates of RBD-binding titers represented in A.

**Figure S3. Related to Figure 3. Cellular response characterization in mice**. (A-B) Spot forming units (SFU) detected by IFN-γ (D) or IL-4 (E) ELISpot in splenocytes extracted from BALB/c animals 4 weeks after vaccination with 10^11^ gc of AC1 or AC3 and stimulated with 2 μg/ml concanavalin A (positive control) for 48h. (C-D) Spot forming units (SFU) detected by IFN-γ (A) or IL-4 (B) ELISpot in splenocytes extracted from C57BL/6 animals 6 weeks after vaccination with 10^10^ gc of AC1 or AC3 and stimulated with 2 μg/ml concanavalin A (positive control) for 48h.

**Figure S4. Related to Figures 5 and 6. Serum chemistry and complete blood counts in NHP**. Measurement of total protein (mg/dL), albumin (g/dL), globulin (g/dL), albumin/globulin ratio (A/G ratio), aspartate aminotransferase (AST or SGOT, measured in IU/L), alanine aminotransferase (ALT or SGPT, measured in IU/L), alkaline phosphatase (ALP, measured in IU/L), gamma-glutamyltransferase (GGT, measured in IU/L), total bilirubin (mg/dL), blood urea (BUN, measured in mg/dL), creatinine (mg/dL), BUN/creatinine ratio, phosphorus (mg/dL), glucose (mg/dL), calcium (mg/dL), magnesium (mEq/dL), sodium (mEq/dL), potassium (mEq/dL), sodium to potassium ratio (Na/K ratio), chloride (mEq/dL), cholesterol (mg/dL), triglycerides (mg/dL), amylase (IU/L), creatine phosphokinase (CPK, measured in IU/L) in serum samples of NHP treated with 10^12^ gc of AC1 or AC3 7 days before injection, on the day of injection and 1, 7 and 14 days post-treatment. Hematological analysis of white blood cells (WBC, ×10^3^cells/µL), red blood cells (RBC, ×10^6^cells/µL), hemoglobin (HB, g/dL), hematocrit (HCT, %), mean corpuscular volume (MCV, fL), mean corpuscular hemoglobin (MCH, pg), mean corpuscular hemoglobin concentration (MCHC, g/dL), platelets (×10^3^cells/µL), absolute neutrophils (cells/µL), absolute lymphocytes (cells/µL), absolute monocytes (cells/µL), absolute eosinophils (cells/µL), prothrombin (seconds), activated partial thromboplastin time (APTT, measured in seconds), fibrinogen (mg/dL) and D-dimers (ng/mL) in EDTA-blood samples of NHP treated with 10^12^ gc of AC1 or AC3 7 days before injection, on the day of injection and 1, 7 and 14 days post-treatment.

**Figure S5. Related to Figures 5 and 6. Serum Cytokine response to AC1 and AC3 in NHP**. Concentration (pg/mL) of IL10, IL2, IL12/23, IL1Rα, IL13, granulocyte-macrophage colony-stimulating factor (GM-CSF, MCP1), IL15, vascular endothelial growth factor (VEGF), IL17α, IL18, transforming growth factor alpha (TGFα) and IL8 in serum samples of NHP treated with 10^12^ gc of AC1 or AC3 7 days before injection, on the day of injection and 1, 7, 14, 21 and 28 days post-treatment.

**Figure S6. Related to Figure 6. Analysis of AAVCOVID SARS-CoV-2 Spike specific T-cell response in NHP**. (A) Schematic representation of portion of the Spike protein represented in each peptide pool used for NHP PBMC stimulation. (B) Flow cytometry scatter plots from AC3 female animal showing the frequency of CD107^+^IFNγ^+^ cells within blood Memory CD8^+^ T cells at baseline and at weeks 6 and 14 post-vaccination. The numbers indicate the frequency within the parent population. (C) Flow cytometry scatter plots from AC3 female animal showing the frequency of TNFα^+^ IL2^+^ cells within blood Memory CD4^+^ T cells at baseline and at weeks 6 and 14 post-vaccination. The numbers indicate the frequency within the parent population.

**Figure S7. Related to Figures 5 and 6. Cellular and humoral anti-AAV capsid response in NHP**. (A) Neutralizing antibody titers against the injected vector (AAVrh32.33) and cross-reactive neutralizing against other serotypes (AAV1, AAV2, AAV5, AAV8, AAV9). (B) Quantification of spot forming units (SFU) by ELISpot in PBMC samples collected at different timepoints in animals treated with AC1 or AC3 and stimulated with peptides spanning AAVrh32.33 capsid sequence.

**Figure S8. Related to Figure 2. Kinetics of biodistribution and expression in mice**. (A) Quantification of vector genome copies (DNA) in the right gastrocnemius (right gastroc) or injection site, left gastrocnemius (left gastroc) or contralateral muscle, liver and spleen on weeks 1, 4 and 8 after the administration of 10^11^ gc of AC1 or AC3 in C57BL/6 animals (n=6, 3/gender). Horizontal dotted lines indicate background levels for right and left gastrocnemius muscles. Liver and spleen had not detectable background. Data are represented as mean±SD. (B) Quantification of transgene expression in the right gastrocnemius muscle on weeks 1, 4 and 8 in C57BL/6 animals (n=3/gender) injected with 10^11^ gc of AC1 or AC3. Data are represented as mean±SD. (C) Table of transgene expression values in experiment describe in A. ND: not detected. (D) Biodistribution of AC3 in several organs 8 weeks after vaccination of BALB/c females treated with 10^11^ gc. Data are represented as mean±SD.

## BIBLIOGRAPHY

1. Anderson, E.J., Rouphael, N.G., Widge, A.T., Jackson, L.A., Roberts, P.C., Makhene, M., Chappell, J.D., Denison, M.R., Stevens, L.J., Pruijssers, A.J., et al. (2020). Safety and Immunogenicity of SARS-CoV-2 mRNA-1273 Vaccine in Older Adults. N Engl J Med.

2. Bennett, J., Wellman, J., Marshall, K., McCague, S., Ashtari, M., DiStefano-Pappas, J., Elci, O., Chung, D., Sun, J., Wright, J., et al. (2016). Safety and durability of effect of contralateral-eye administration of AAV2 gene therapy in patients with childhood-onset blindness caused by RPE65 mutations: a follow-on phase 1 trial. Lancet (London, England) 388.

3. Biswas, M., Rahaman, S., Biswas, T., Haque, Z., and Ibrahim, B. (2020). Association of Sex, Age, and Comorbidities with Mortality in COVID-19 Patients: A Systematic Review and Meta-Analysis. Intervirology.

4. Calcedo, R., Chichester, J., and Wilson, J. (2018). Assessment of Humoral, Innate, and T-Cell Immune Responses to Adeno-Associated Virus Vectors. Human gene therapy methods 29.

5. Calcedo, R., Vandenberghe, L.H., Gao, G., Lin, J., and Wilson, J.M. (2009). Worldwide epidemiology of neutralizing antibodies to adeno-associated viruses. J Infect Dis 199, 381–390.

6. Case, J., Rothlauf, P., Chen, R., Kafai, N., Fox, J., Smith, B., Shrihari, S., McCune, B., Harvey, IA, Keeler, S., Bloyet, L., et al. (2020). Replication-Competent Vesicular Stomatitis Virus Vaccine Vector Protects against SARS-CoV-2-Mediated Pathogenesis in Mice (CellPress: Cell Host & Microbe).

7. Chand, D., Mohr, F., McMillan, H., Tukov, F.F., Montgomery, K., Kleyn, A., Sun, R., Tauscher-Wisniewski, S., Kaufmann, P., and Kullak-Ublick, G. (2020). Hepatotoxicity following administration of onasemnogene abeparvovec (AVXS-101) for the treatment of spinal muscular atrophy. J Hepatol.

8. Chandrashekar, A., Liu, J., Martinot, A.J., McMahan, K., Mercado, N.B., Peter, L., Tostanoski, L.H., Yu, J., Maliga, Z., Nekorchuk, M., et al. (2020). SARS-CoV-2 infection protects against rechallenge in rhesus macaques. Science 369, 812–817.

9. Corbett, K., Flynn, B., Foulds, K., Francica, J., Boyoglu-Barnum, S., Werner, A., Flach, B., O’Connell, S., Bock, K., Minai, M., et al. (2020). Evaluation of the mRNA-1273 Vaccine against SARS-CoV-2 in Nonhuman Primates. The New England journal of medicine 383.

10. Crawford, K., Eguia, R., Dingens, A., Loes, A., Malone, K., Wolf, C., Chu, H., Tortorici, M., Veesler, D., Murphy, M., et al. (2020). Protocol and Reagents for Pseudotyping Lentiviral Particles with SARS-CoV-2 Spike Protein for Neutralization Assays. Viruses 12.

11. Cuschieri, S., and Grech, S. (2020). Obesity population at risk of COVID-19 complications. Global health, epidemiology and genomics 5.

12. Darriba, D., Taboada, G.L., Doallo, R., and Posada, D. (2011). ProtTest 3: fast selection of best-fit models of protein evolution. Bioinformatics 27, 1164–1165.

13. Fausther-Bovendo, H., and Kobinger, G.P. (2014). Pre-existing immunity against Ad vectors: humoral, cellular, and innate response, what’s important? Hum Vaccin Immunother 10, 2875–2884.

14. Feng, L., Wang, Q., Shan, C., Yang, C., Feng, Y., Wu, J., Liu, X., Zhou, Y., Jiang, R., Hu, P., et al. (2020). An adenovirus-vectored COVID-19 vaccine confers protection from SARS-COV-2 challenge in rhesus macaques. Nature communications 11.

15. Folegatti, P., Bittaye, M., Flaxman, A., Lopez, F., Bellamy, D., Kupke, A., Mair, C., Makinson, R., Sheridan, J., Rohde, C., et al. (2020a). Safety and immunogenicity of a candidate Middle East respiratory syndrome coronavirus viral-vectored vaccine: a dose-escalation, open-label, non-randomised, uncontrolled, phase 1 trial. The Lancet Infectious diseases 20.

16. Folegatti, P., Ewer, K., Aley, P., Angus, B., Becker, S., Belij-Rammerstorfer, S., Bellamy, D., Bibi, S., Bittaye, M., Clutterbuck, E., et al. (2020b). Safety and immunogenicity of the ChAdOx1 nCoV-19 vaccine against SARS-CoV-2: a preliminary report of a phase 1/2, single-blind, randomised controlled trial. Lancet (London, England) 396.

17. Gao, G., Vandenberghe, L.H., Alvira, M.R., Lu, Y., Calcedo, R., Zhou, X., and Wilson, J.M. (2004). Clades of Adeno-associated viruses are widely disseminated in human tissues. J Virol 78, 6381–6388.

18. Gao, G., Wang, Q., Calcedo, R., Mays, L., Bell, P., Wang, L., Vandenberghe, L.H., Grant, R., Sanmiguel, J., Furth, E.E., et al. (2009). Adeno-associated virus-mediated gene transfer to nonhuman primate liver can elicit destructive transgene-specific T cell responses. Hum Gene Ther 20, 930–942.

19. Graham, S.P., McLean, R.K., Spencer, A.J., Belij-Rammerstorfer, S., Wright, D., Ulaszewska, M., Edwards, J.C., Hayes, J.W.P., Martini, V., Thakur, N., et al. (2020). Evaluation of the immunogenicity of prime-boost vaccination with the replication-deficient viral vectored COVID-19 vaccine candidate ChAdOx1 nCoV-19. NPJ Vaccines 5, 69.

20. Greig, J.A., Calcedo, R., Grant, R.L., Peng, H., Medina-Jaszek, C.A., Ahonkhai, O., Qin, Q., Roy, S., Tretiakova, A.P., and Wilson, J.M. (2016). Intramuscular administration of AAV overcomes pre-existing neutralizing antibodies in rhesus macaques. Vaccine 34, 6323–6329.

21. Guindon, S., and Gascuel, O. (2003). A simple, fast, and accurate algorithm to estimate large phylogenies by maximum likelihood. Syst Biol 52, 696–704.

22. Hassan, A., Kafai, N., Dmitriev, I., Fox, J., Smith, B., Harvey, I., Chen, R., Winkler, E., Wessel, A., Case, J., et al. (2020). A Single-Dose Intranasal ChAd Vaccine Protects Upper and Lower Respiratory Tracts against SARS-CoV-2. Cell 183.

23. High, K.A., and Roncarolo, M.G. (2019). Gene Therapy. N Engl J Med 381, 455–464.

24. Jackson, L., Anderson, E., Rouphael, N., Roberts, P., Makhene, M., Coler, R., McCullough, M., Chappell, J., Denison, M., Stevens, L., et al. (2020). An mRNA Vaccine against SARS-CoV-2 - Preliminary Report. The New England journal of medicine 383.

25. Johnson, J.L., Rosenthal, R.L., Knox, J.J., Myles, A., Naradikian, M.S., Madej, J., Kostiv, M., Rosenfeld, A.M., Meng, W., Christensen, S.R., et al. (2020). The Transcription Factor T-bet Resolves Memory B Cell Subsets with Distinct Tissue Distributions and Antibody Specificities in Mice and Humans. Immunity 52, 842–855 e846.

26. Kalnin, K.V., Plitnik, T., Kishko, M., Zhang, J., Zhang, D., Beauvais, A., Anosova, N.G., Tibbitts, T., DiNapoli, J.M., Huang, P.-W.D., et al. (2020). Immunogenicity of novel mRNA COVID-19 vaccine MRT5500 in mice and non-human primates.

27. Karlsson, E., Hertz, T., Johnson, C., Mehle, A., Krammer, F., and Schultz-Cherry, S. (2016). Obesity Outweighs Protection Conferred by Adjuvanted Influenza Vaccination. mBio 7.

28. Kim, Y.-H., Center, A.M., Kim, J.-K., Viral Infectious Disease Research Center, K.R.I.o.B., Biotechnology, D., Kim, D.-J., Viral Infectious Disease Research Center, K.R.I.o.B., Biotechnology, D., Nam, J.-H., Viral Infectious Disease Research Center, K.R.I.o.B., et al. (2020). Diet-Induced Obesity Dramatically Reduces the Efficacy of a 2009 Pandemic H1N1 Vaccine in a Mouse Model. The Journal of Infectious Diseases 205, 244-251.

29. Knox, J.J., Buggert, M., Kardava, L., Seaton, K.E., Eller, M.A., Canaday, D.H., Robb, M.L., Ostrowski, M.A., Deeks, S.G., Slifka, M.K., et al. (2017). T-bet+ B cells are induced by human viral infections and dominate the HIV gp140 response. JCI Insight 2.

30. Koch, T., Dahlke, C., Fathi, A., Kupke, A., Krähling, V., Okba, N., Halwe, S., Rohde, C., Eickmann, M., Volz, A., et al. (2020). Safety and immunogenicity of a modified vaccinia virus Ankara vector vaccine candidate for Middle East respiratory syndrome: an open-label, phase 1 trial. The Lancet Infectious diseases 20.

31. Kremsner, P., Mann, P., Bosch, J., Fendel, R., Gabor, J.J., Kreidenweiss, A., Kroidl, A., Leroux-Roels, I., Leroux-Roels, G., Schindler, C., et al. (2020). Phase 1 Assessment of the Safety and Immunogenicity of an mRNA-Lipid Nanoparticle Vaccine Candidate Against SARS-CoV-2 in Human Volunteers. medRxiv.

32. Kumar, S., Stecher, G., Li, M., Knyaz, C., and Tamura, K. (2018). MEGA X: Molecular Evolutionary Genetics Analysis across Computing Platforms. Mol Biol Evol 35, 1547-1549.

33. Kurtz, S., Choudhuri, J., Ohlebusch, E., Schleiermacher, C., Stoye, J., and Giegerich, R. (2001). REPuter: the manifold applications of repeat analysis on a genomic scale. Nucleic acids research 29.

34. Kutyavin, I., Afonina, I., Mills, A., Gorn, V., Lukhtanov, E., Belousov, E., Singer, M., Walburger, D., Lokhov, S., Gall, A., et al. (2000). 3’-minor groove binder-DNA probes increase sequence specificity at PCR extension temperatures. Nucleic acids research 28.

35. Le, S.Q., and Gascuel, O. (2008). An improved general amino acid replacement matrix. Mol Biol Evol 25, 1307–1320.

36. Li, X., Cao, H., Wang, Q., Di, B., Wang, M., Lu, J., Pan, L., Yang, L., Mei, M., Pan, X., et al. (2012). Novel AAV-based genetic vaccines encoding truncated dengue virus envelope proteins elicit humoral immune responses in mice. Microbes Infect 14, 1000–1007.

37. Lin, J., Calcedo, R., Vandenberghe, L.H., Bell, P., Somanathan, S., and Wilson, J.M. (2009). A new genetic vaccine platform based on an adeno-associated virus isolated from a rhesus macaque. J Virol 83, 12738–12750.

38. Livak, K., and Schmittgen, T. (2001). Analysis of relative gene expression data using real-time quantitative PCR and the 2(-Delta Delta C(T)) Method. Methods (San Diego, Calif) 25.

39. Lock, M., Alvira, M., Vandenberghe, L.H., Samanta, A., Toelen, J., Debyser, Z., and Wilson, J.M. (2010). Rapid, simple, and versatile manufacturing of recombinant adeno-associated viral vectors at scale. Hum Gene Ther 21, 1259–1271.

40. Logunov, D., Dolzhikova, I., Zubkova, O., Tukhvatullin, A., Shcheblyakov, D., Dzharullaeva, A., Grousova, D., Erokhova, A., Kovyrshina, A., Botikov, A., et al. (2020). Safety and immunogenicity of an rAd26 and rAd5 vector-based heterologous prime-boost COVID-19 vaccine in two formulations: two open, non-randomised phase 1/2 studies from Russia. Lancet (London, England) 396.

41. Majowicz, A., Salas, D., Zabaleta, N., Rodríguez-Garcia, E., González-Aseguinolaza, G., Petry, H., and Ferreira, V. (2017). Successful Repeated Hepatic Gene Delivery in Mice and Non-human Primates Achieved by Sequential Administration of AAV5 ch and AAV1. Molecular therapy: the journal of the American Society of Gene Therapy 25.

42. Markham, N., and Zuker, M. (2005). DINAMelt web server for nucleic acid melting prediction. Nucleic acids research 33.

43. Markham, N., and Zuker, M. (2008). UNAFold: software for nucleic acid folding and hybridization. Methods in molecular biology (Clifton, NJ) 453.

44. Martin, J., Louder, M., Holman, L., Gordon, I., Enama, M., Larkin, B., Andrews, C., Vogel, L., Koup, R., Roederer, M., et al. (2008). A SARS DNA vaccine induces neutralizing antibody and cellular immune responses in healthy adults in a Phase I clinical trial. Vaccine 26.

45. Mays, L.E., Vandenberghe, L.H., Xiao, R., Bell, P., Nam, H.J., Agbandje-McKenna, M., and Wilson, J.M. (2009). Adeno-associated virus capsid structure drives CD4-dependent CD8+ T cell response to vector encoded proteins. J Immunol 182, 6051–6060.

46. Mays, L.E., Wang, L., Tenney, R., Bell, P., Nam, H.J., Lin, J., Gurda, B., Van Vliet, K., Mikals, K., Agbandje-McKenna, M., et al. (2013). Mapping the structural determinants responsible for enhanced T cell activation to the immunogenic adeno-associated virus capsid from isolate rhesus 32.33. J Virol 87, 9473–9485.

47. McMahan, K., Yu, J., Mercado, N.B., Loos, C., Tostanoski, L.H., Chandrashekar, A., Liu, J., Peter, L., Atyeo, C., Zhu, A., et al. (2020). Correlates of protection against SARS-CoV-2 in rhesus macaques. Nature, 1-8.

48. Mehendale, S., van Lunzen, J., Clumeck, N., Rockstroh, J., Vets, E., Johnson, P.R., Anklesaria, P., Barin, B., Boaz, M., Kochhar, S., et al. (2008). A phase 1 study to evaluate the safety and immunogenicity of a recombinant HIV type 1 subtype C adeno-associated virus vaccine. AIDS Res Hum Retroviruses 24, 873–880.

49. Mendell, J., Al-Zaidy, S., Shell, R., Arnold, W., Rodino-Klapac, L., Prior, T., Lowes, L., Alfano, L., Berry, K., Church, K., et al. (2017). Single-Dose Gene-Replacement Therapy for Spinal Muscular Atrophy. The New England journal of medicine 377.

50. Mercado, N.B., Zahn, R., Wegmann, F., Loos, C., Chandrashekar, A., Yu, J., Liu, J., Peter, L., McMahan, K., Tostanoski, L.H., et al. (2020). Single-shot Ad26 vaccine protects against SARS-CoV-2 in rhesus macaques. Nature 586, 583–588.

51. Mingozzi, F., and High, K.A. (2017). Overcoming the Host Immune Response to Adeno-Associated Virus Gene Delivery Vectors: The Race Between Clearance, Tolerance, Neutralization, and Escape. Annu Rev Virol 4, 511–534.

52. Modjarrad, K., Roberts, C., Mills, K., Castellano, A., Paolino, K., Muthumani, K., Reuschel, E., Robb, M., Racine, T., Oh, M., et al. (2019). Safety and immunogenicity of an anti-Middle East respiratory syndrome coronavirus DNA vaccine: a phase 1, open-label, single-arm, dose-escalation trial. The Lancet Infectious diseases 19.

53. Muthumani, K., Falzarano, D., Reuschel, E., Tingey, C., Flingai, S., Villarreal, D., Wise, M., Patel, A., Izmirly, A., Aljuaid, A., et al. (2015). A synthetic consensus anti-spike protein DNA vaccine induces protective immunity against Middle East respiratory syndrome coronavirus in nonhuman primates. Science translational medicine 7.

54. Nieto, K., and Salvetti, A. (2014). AAV Vectors Vaccines Against Infectious Diseases. Front Immunol 5, 5.

55. Pacouret, S., Bouzelha, M., Shelke, R., Andres-Mateos, E., Xiao, R., Maurer, A., Mevel, M., Turunen, H., Barungi, T., Penaud-Budloo, M., et al. (2017). AAV-ID: A Rapid and Robust Assay for Batch-to-Batch Consistency Evaluation of AAV Preparations. Molecular therapy: the journal of the American Society of Gene Therapy 25.

56. Park, H., Shim, S., Lee, E., Cho, W., Park, S., Jeon, H., Ahn, S., Kim, H., and JH, N. (2014). Obesity-induced chronic inflammation is associated with the reduced efficacy of influenza vaccine (Human Vaccines and Immunotherapeutics), pp. 1181–1186.

57. Patel, A., Walters, J., Reuschel, E.L., Schultheis, K., Parzych, E., Gary, E.N., Maricic, I., Purwar, M., Eblimit, Z., Walker, S.N., et al. (2020). Intradermal-delivered DNA vaccine provides anamnestic protection in a rhesus macaque SARS-CoV-2 challenge model.

58. R, C., LH, V., G, G., J, L., and JM, W. (2009). Worldwide epidemiology of neutralizing antibodies to adeno-associated viruses. The Journal of infectious diseases 199.

59. Salganik, M., Hirsch, M.L., and Samulski, R.J. (2015). Adeno-associated Virus as a Mammalian DNA Vector. Microbiology Spectrum.

60. Sanchez-Felipe, L., Vercruysse, T., Sharma, S., Ma, J., Lemmens, V., Looveren, D.V., Javarappa, M.P.A., Boudewijns, R., Malengier-Devlies, B., Liesenborghs, L., et al. (2020). A single-dose live-attenuated YF17D-vectored SARS-CoV-2 vaccine candidate. Nature, 1-10.

61. Sanmiguel, J., Gao, G., and Vandenberghe, L. (2019). Quantitative and Digital Droplet-Based AAV Genome Titration. Methods in molecular biology (Clifton, NJ) 1950.

62. Sievers, F., and Higgins, D.G. (2018). Clustal Omega for making accurate alignments of many protein sequences. Protein Sci 27, 135–145.

63. Stroes, E., Nierman, M., Meulenberg, J., Franssen, R., Twisk, J., Henny, C., Maas, M., Zwinderman, A., Ross, C., Aronica, E., et al. (2008). Intramuscular administration of AAV1-lipoprotein lipase S447X lowers triglycerides in lipoprotein lipase-deficient patients. Arteriosclerosis, thrombosis, and vascular biology 28.

64. van Doremalen, N., Haddock, E., Feldmann, F., Meade-White, K., Bushmaker, T., Fischer, R., Okumura, A., Hanley, P., Saturday, G., Edwards, N., et al. (2020a). A single dose of ChAdOx1 MERS provides protective immunity in rhesus macaques. Science advances 6.

65. van Doremalen, N., Lambe, T., Spencer, A., Belij-Rammerstorfer, S., Purushotham, J., Port, J., Avanzato, V., Bushmaker, T., Flaxman, A., Ulaszewska, M., et al. (2020b). ChAdOx1 nCoV-19 vaccine prevents SARS-CoV-2 pneumonia in rhesus macaques. Nature 586.

66. Vandenberghe, L.H., Breous, E., Nam, H.J., Gao, G., Xiao, R., Sandhu, A., Johnston, J., Debyser, Z., Agbandje-McKenna, M., and Wilson, J.M. (2009a). Naturally occurring singleton residues in AAV capsid impact vector performance and illustrate structural constraints. Gene Ther 16, 1416–1428.

67. Vandenberghe, L.H., and Wilson, J.M. (2007). AAV as an immunogen. Curr Gene Ther 7, 325–333.

68. Vandenberghe, L.H., Wilson, J.M., and Gao, G. (2009b). Tailoring the AAV vector capsid for gene therapy. Gene Ther 16, 311–319.

69. Vardas, E., Kaleebu, P., Bekker, L.G., Hoosen, A., Chomba, E., Johnson, P.R., Anklesaria, P., Birungi, J., Barin, B., Boaz, M., et al. (2010). A phase 2 study to evaluate the safety and immunogenicity of a recombinant HIV type 1 vaccine based on adeno-associated virus. AIDS Res Hum Retroviruses 26, 933–942.

70. Walls, A., Park, Y., Tortorici, M., Wall, A., McGuire, A., and Veesler, D. (2020). Structure, Function, and Antigenicity of the SARS-CoV-2 Spike Glycoprotein. Cell 181.

71. Walsh, E., Frenck, R., Falsey, A., Kitchin, N., Absalon, J., Gurtman, A., Lockhart, S., Neuzil, K., Mulligan, M., Bailey, R., et al. (2020). Safety and Immunogenicity of Two RNA-Based Covid-19 Vaccine Candidates. The New England journal of medicine.

72. Wang, Q., Zhang, Y., Wu, L., Niu, S., Song, C., Zhang, Z., Lu, G., Qiao, C., Hu, Y., Yuen, K., et al. (2020). Structural and Functional Basis of SARS-CoV-2 Entry by Using Human ACE2. Cell 181.

73. Wang, X., Yan, Y., Gan, T., Yang, X., Li, D., Zhou, D., Sun, Q., Huang, Z., and Zhong, J. (2019). A trivalent HCV vaccine elicits broad and synergistic polyclonal antibody response in mice and rhesus monkey. Gut 68.

74. Widge, A.T., Rouphael, N.G., Jackson, L.A., Anderson, E.J., Roberts, P.C., Makhene, M., Chappell, J.D., Denison, M.R., Stevens, L.J., Pruijssers, A.J., et al. (2020). Durability of Responses after SARS-CoV-2 mRNA-1273 Vaccination. N Engl J Med.

75. Wrapp, D., Wang, N., Corbett, K., Goldsmith, J., Hsieh, C., Abiona, O., Graham, B., and McLellan, J. (2020). Cryo-EM structure of the 2019-nCoV spike in the prefusion conformation. Science (New York, NY) 367.

76. Wu, F., Zhao, S., Yu, B., Chen, Y.-M., Wang, W., Song, Z.-G., Hu, Y., Tao, Z.-W., Tian, J.-H., Pei, Y.-Y., et al. (2020). A new coronavirus associated with human respiratory disease in China. Nature 579, 265–269.

77. Yu, J., Tostanoski, L.H., Peter, L., Mercado, N.B., McMahan, K., Mahrokhian, S.H., Nkolola, J.P., Liu, J., Li, Z., Chandrashekar, A., et al. (2020). DNA vaccine protection against SARS-CoV-2 in rhesus macaques. Science 369, 806–811.

78. Zhang, N., Li, X., Deng, Y., Zhao, H., Huang, Y., Yang, G., Huang, W., Gao, P., Zhou, C., Zhang, R., et al. (2020). A Thermostable mRNA Vaccine against COVID-19. Cell 182.

79. Zhou, P., Yang, X., Wang, X., Hu, B., Zhang, L., Zhang, W., Si, H., Zhu, Y., Li, B., Huang, C., et al. (2020). A pneumonia outbreak associated with a new coronavirus of probable bat origin. Nature 579.

80. Zhu, F., Chen, T., Zhang, Y., Sun, H., Cao, H., Lu, J., Zhao, L., and Li, G. (2015). A Novel Adeno-Associated Virus-Based Genetic Vaccine Encoding the Hepatitis C Virus NS3/4 Protein Exhibits Immunogenic Properties in Mice Superior to Those of an NS3-Protein-Based Vaccine. PLoS One 10, e0142349.

81. Zhu, F., Guan, X., Li, Y., Huang, J., Jiang, T., Hou, L., Li, J., Yang, B., Wang, L., Wang, W., et al. (2020a). Immunogenicity and safety of a recombinant adenovirus type-5-vectored COVID-19 vaccine in healthy adults aged 18 years or older: a randomised, double-blind, placebo-controlled, phase 2 trial. Lancet (London, England) 396.

82. Zhu, F., Li, Y., Guan, X., Hou, L., Wang, W., Li, J., Wu, S., Wang, B., Wang, Z., Wang, L., et al. (2020b). Safety, tolerability, and immunogenicity of a recombinant adenovirus type-5 vectored COVID-19 vaccine: a dose-escalation, open-label, non-randomised, first-in-human trial. Lancet (London, England) 395.

83. Zhu, F., Wang, Y., Xu, Z., Qu, H., Zhang, H., Niu, L., Xue, H., Jing, D., and He, H. (2019). Novel adenoassociated virusbased genetic vaccines encoding hepatitis C virus E2 glycoprotein elicit humoral immune responses in mice. Mol Med Rep 19, 1016–1023.

84. Zuker, M. (2003). Mfold web server for nucleic acid folding and hybridization prediction. Nucleic acids research 31.

