## Supplemental Files for "Immunogenicity of an AAV-based, room-temperature stable, single dose COVID-19 vaccine in mouse and non-human primates"

**Figure S8. Related to Figure 2. Kinetics of biodistribution and expression in mice.**

(A) Quantification of vector genome copies (DNA) in the right gastrocnemius (right gastroc) or injection site, left gastrocnemius (left gastroc) or contralateral muscle, liver and spleen on weeks 1, 4 and 8 after the administration of $10^{11}$ gc of AC1 or AC3 in C57BL/6 animals (n=6, 3/gender). Horizontal dotted lines indicate background levels for right and left gastrocnemius muscles. Liver and spleen had not detectable background. Data are represented as mean±SD.

(B) Quantification of transgene expression in the right gastrocnemius muscle on weeks 1, 4 and 8 in C57BL/6 animals (n=3/gender) injected with $10^{11}$ gc of AC1 or AC3. Data are represented as mean±SD.

(C) Table of transgene expression values in experiment describe in A. ND: not detected.

(D) Biodistribution of AC3 in several organs 8 weeks after vaccination of BALB/c females treated with $10^{11}$ gc. Data are represented as mean±SD.

### Figure S1

A

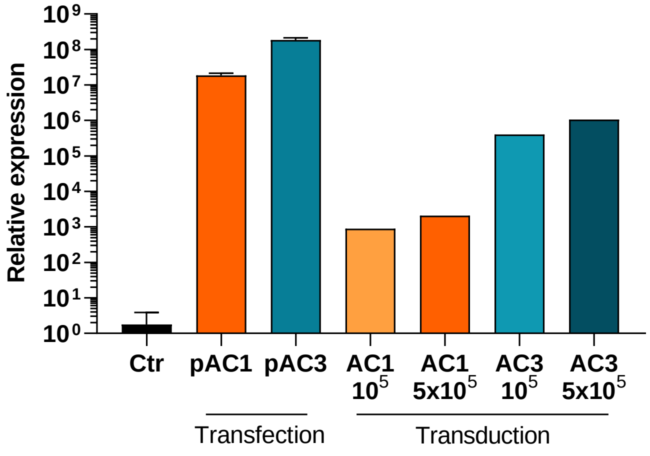

Relative expression
10^9
10^8
10^7
10^6
10^5
10^4
10^3
10^2
10^1
10^0
Ctr
pAC1
pAC3
AC1 10^5
AC1 5x10^5
AC3 10^5
AC3 5x10^5
Transfection
Transduction


B

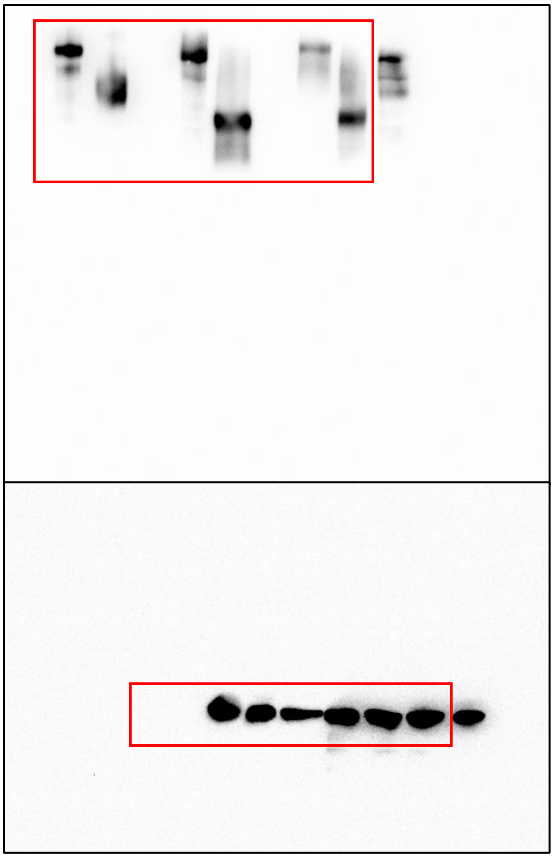

Figure S2

A

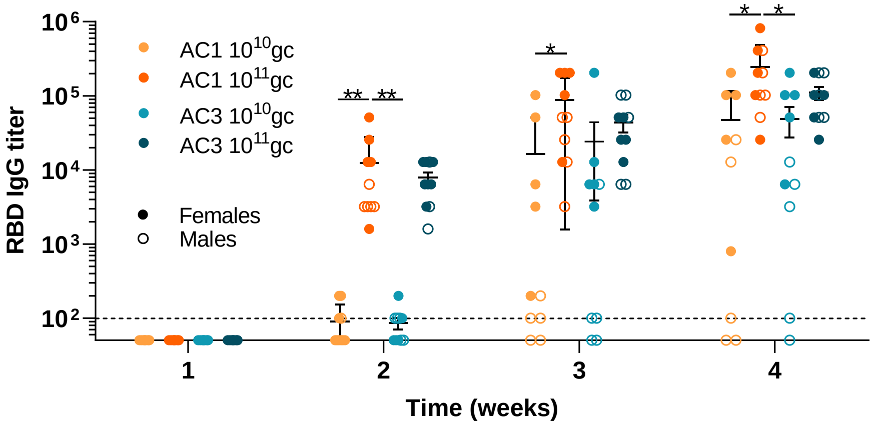

RBD IgG titer
AC1 10^10gc
AC1 10^11gc
AC3 10^10gc
AC3 10^11gc
Females
Males
** **
*
* *
Time (weeks)


B

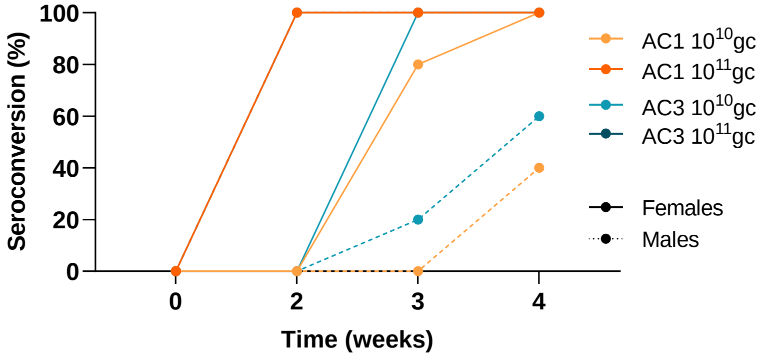

Seroconversion (%)
AC1 10^10gc
AC1 10^11gc
AC3 10^10gc
AC3 10^11gc
Females
Males
Time (weeks)

Figure S3

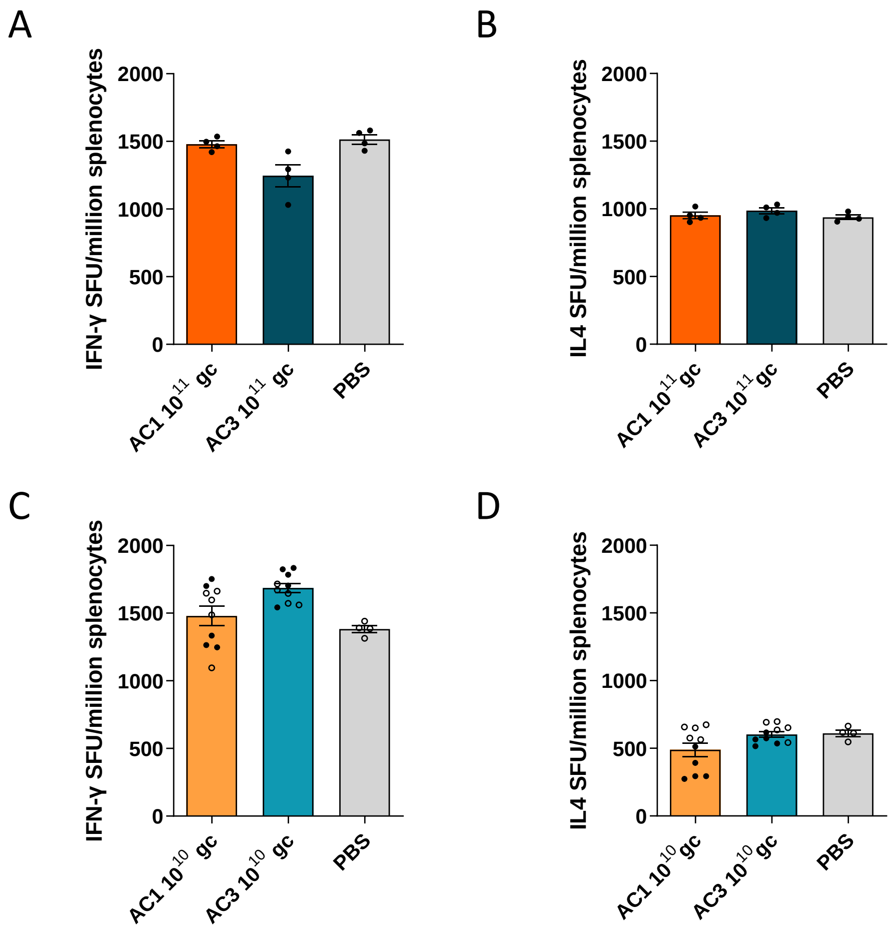

A
IFN-γ SFU/million splenocytes
2000
1500
1000
500
0
AC1 10^11 gc
AC3 10^11 gc
PBS
B
IL4 SFU/million splenocytes
2000
1500
1000
500
0
AC1 10^11 gc
AC3 10^11 gc
PBS
C
IFN-γ SFU/million splenocytes
2000
1500
1000
500
0
AC1 10^10 gc
AC3 10^10 gc
PBS
D
IL4 SFU/million splenocytes
2000
1500
1000
500
0
AC1 10^10 gc
AC3 10^10 gc
PBS

### Figure S4

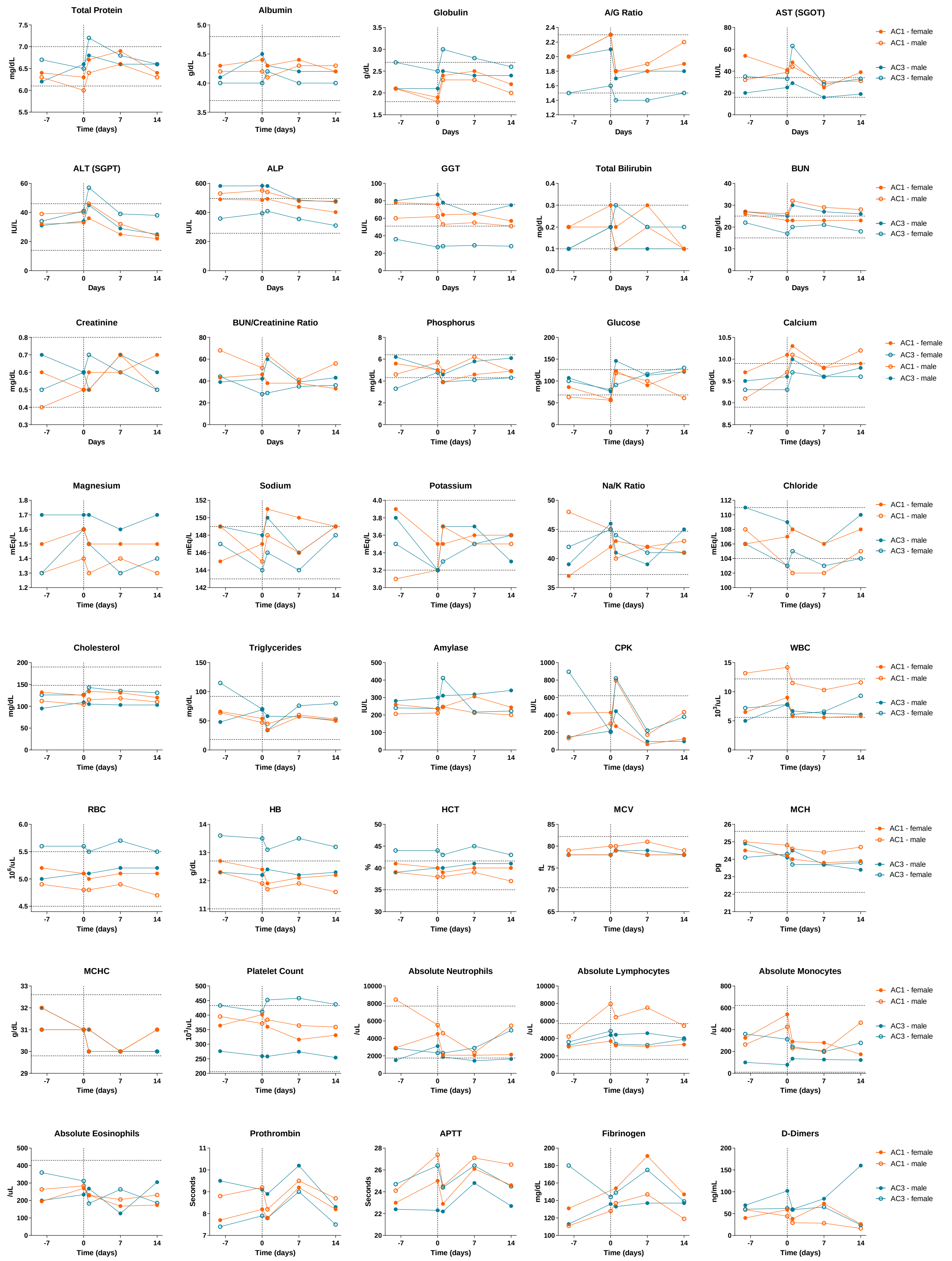
Total Protein
mg/dL
Time (days)
Albumin
g/dL
Time (days)
Globulin
g/dL
Days
A/G Ratio
Days
AST (SGOT)
IU/L
Days
AC1 - female
AC1 - male
AC3 - male
AC3 - female
ALT (SGPT)
IU/L
Days
ALP
IU/L
Days
GGT
IU/L
Days
Total Bilirubin
mg/dL
Days
BUN
mg/dL
Days
AC1 - female
AC1 - male
AC3 - male
AC3 - female
Creatinine
mg/dL
Days
BUN/Creatinine Ratio
Days
Phosphorus
mg/dL
Time (days)
Glucose
mg/dL
Time (days)
Calcium
mg/dL
Time (days)
AC1 - female
AC3 - female
AC1 - male
AC3 - male
Magnesium
mEq/L
Time (days)
Sodium
mEq/L
Time (days)
Potassium
mEq/L
Time (days)
Na/K Ratio
Time (days)
Chloride
mEq/L
Time (days)
AC1 - female
AC1 - male
AC3 - male
AC3 - female
Cholesterol
mg/dL
Time (days)
Triglycerides
mg/dL
Time (days)
Amylase
IU/L
Time (days)
CPK
IU/L
Time (days)
WBC
10³/uL
Time (days)
AC1 - female
AC1 - male
AC3 - male
AC3 - female
RBC
10⁶/uL
Time (days)
HB
g/dL
Time (days)
HCT
%
Time (days)
MCV
fL
Time (days)
MCH
pg
Time (days)
AC1 - female
AC1 - male
AC3 - male
AC3 - female
MCHC
g/dL
Time (days)
Platelet Count
10³/uL
Time (days)
Absolute Neutrophils
/uL
Time (days)
Absolute Lymphocytes
/uL
Time (days)
Absolute Monocytes
/uL
Time (days)
AC1 - female
AC1 - male
AC3 - male
AC3 - female
Absolute Eosinophils
/uL
Time (days)
Prothrombin
Seconds
Time (days)
APTT
Seconds
Time (days)
Fibrinogen
mg/dL
Time (days)
D-Dimers
ng/mL
Time (days)
AC1 - female
AC1 - male
AC3 - male
AC3 - female

### Figure S5

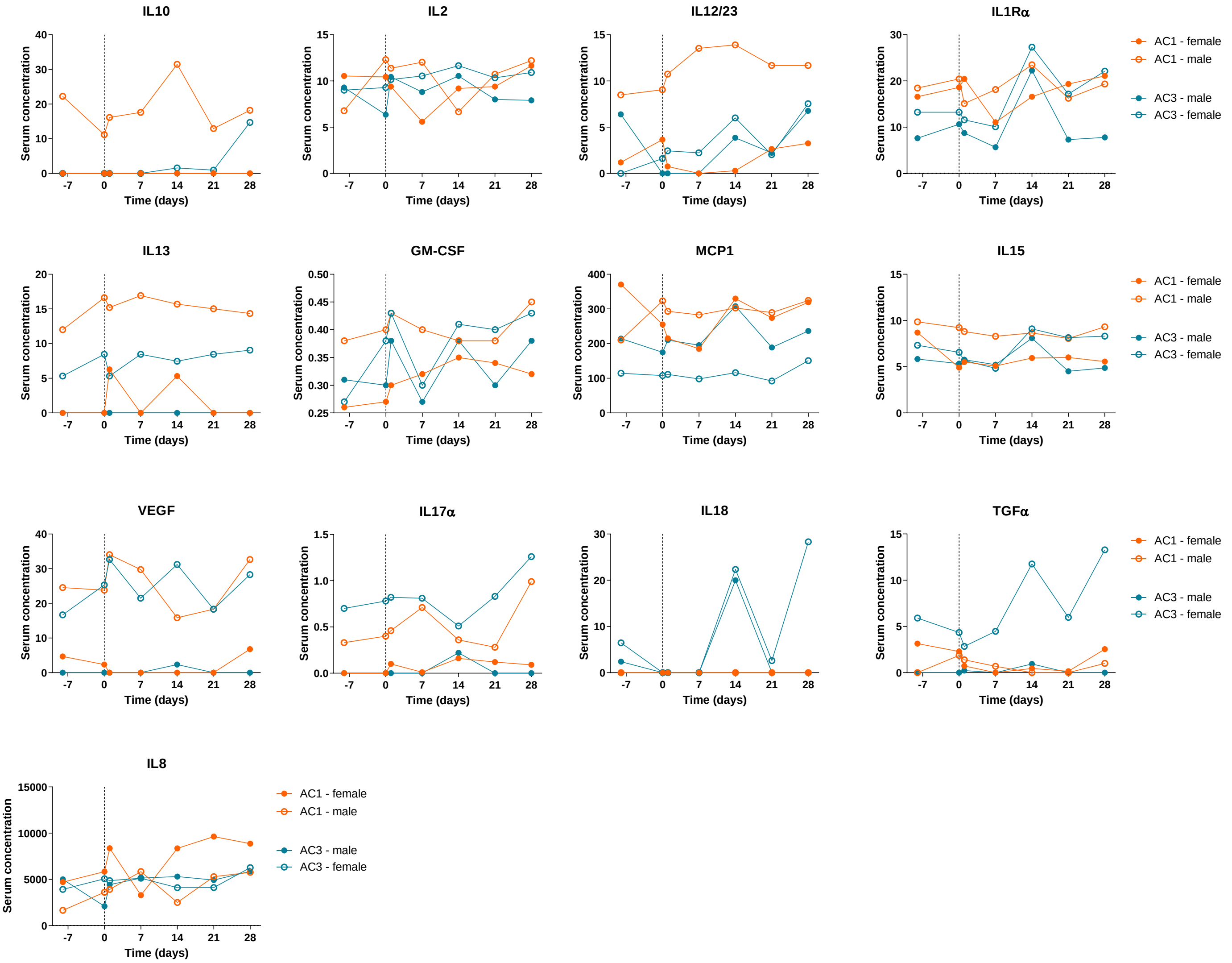
IL10
IL2
IL12/23
IL1Rα
IL13
GM-CSF
MCP1
IL15
VEGF
IL17α
IL18
TGFα
IL8
Serum concentration
Time (days)
-7
0
7
14
21
28
AC1 - female
AC1 - male
AC3 - male
AC3 - female

### Figure S6

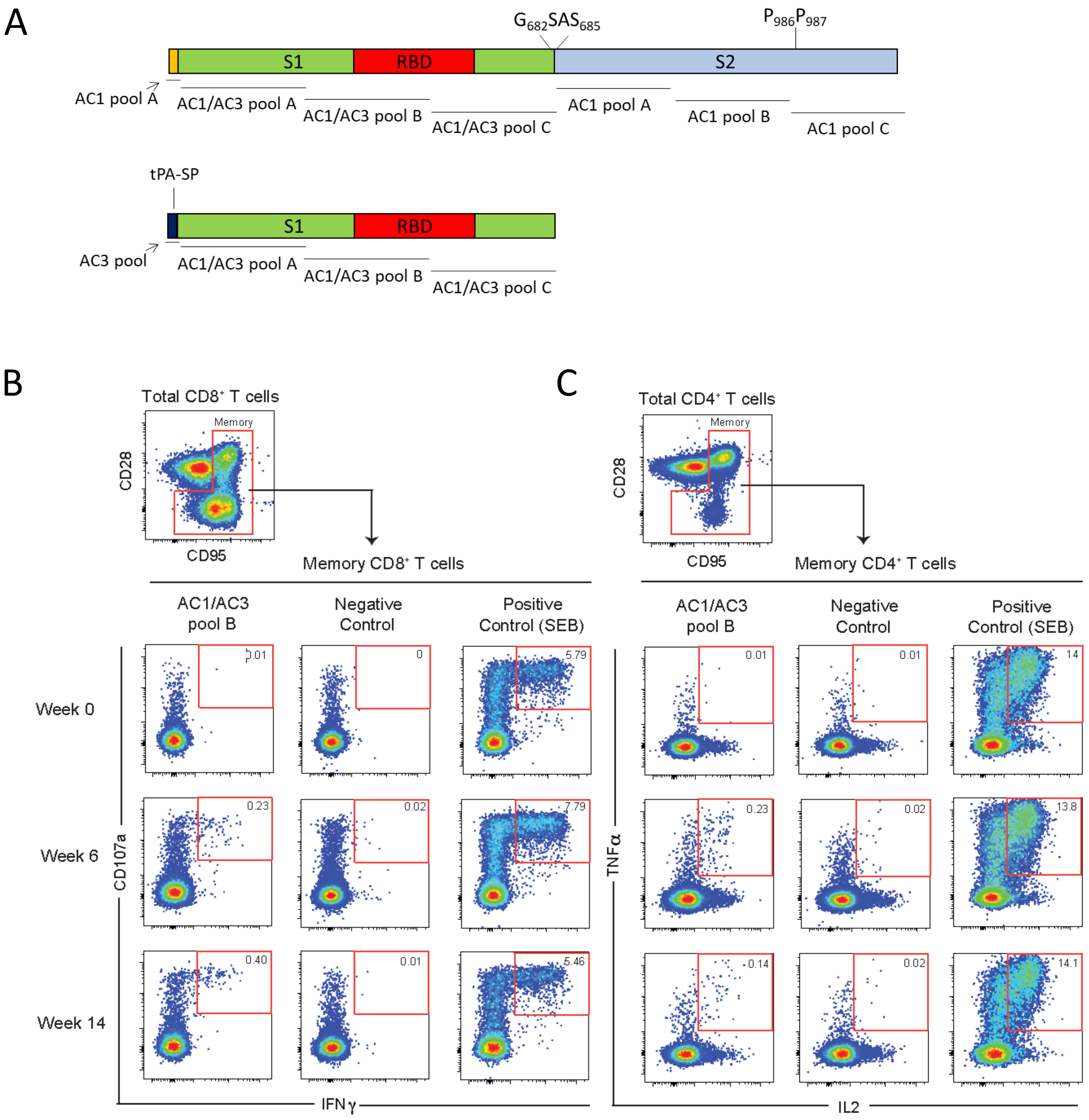
A
$G_{682}SAS_{685}$
$P_{986}P_{987}$
S1
RBD
S2
AC1 pool A
AC1/AC3 pool A
AC1/AC3 pool B
AC1/AC3 pool C
AC1 pool A
AC1 pool B
AC1 pool C
tPA-SP
AC3 pool
B
Total CD8⁺ T cells
Memory
CD28
CD95
Memory CD8⁺ T cells
AC1/AC3 pool B
Negative Control
Positive Control (SEB)
Week 0
Week 6
Week 14
CD107a
IFNγ
C
Total CD4⁺ T cells
Memory CD4⁺ T cells
TNFα
IL2

Figure S7

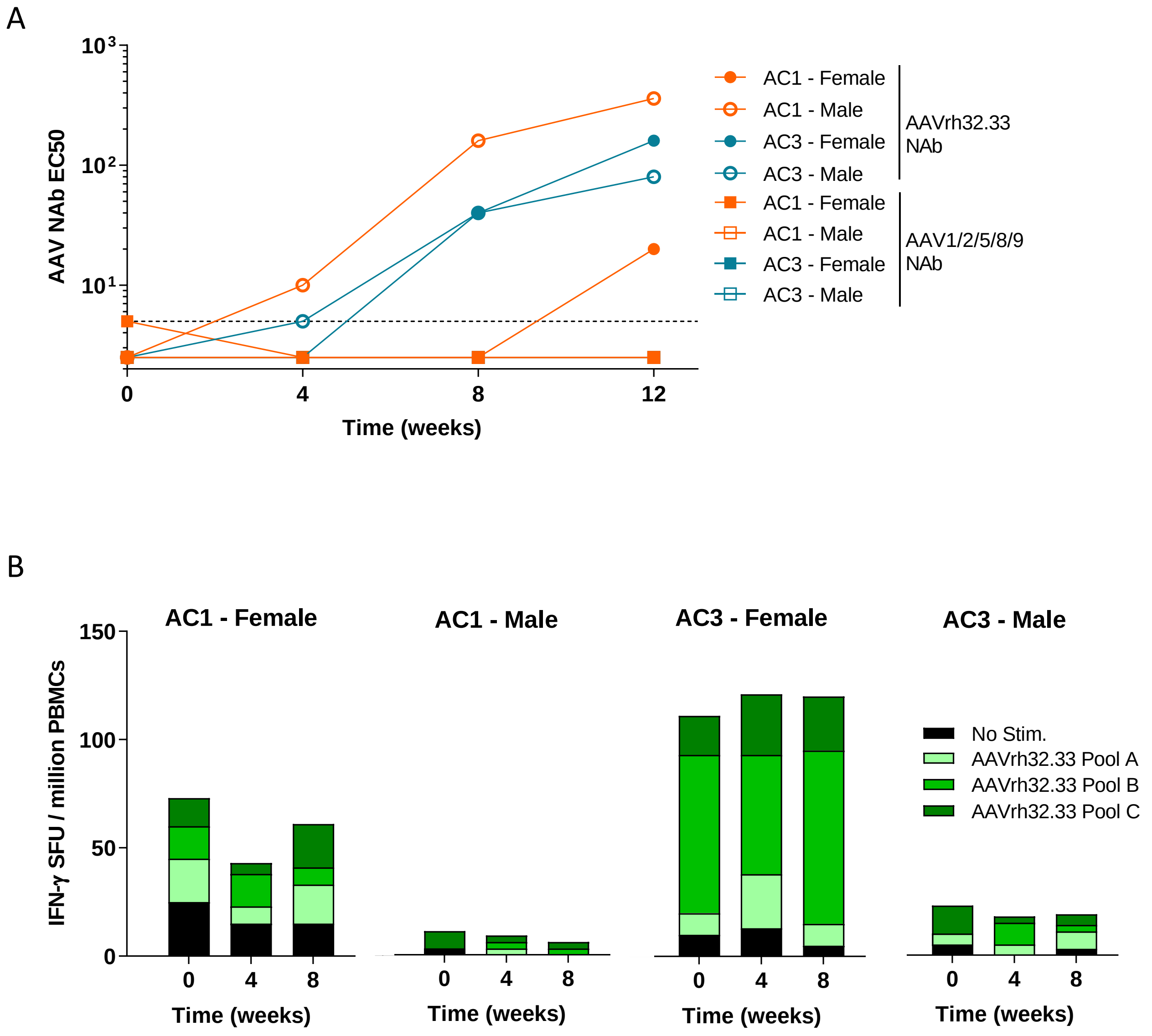
A
AAV NAb EC50
$10^3$
$10^2$
$10^1$
0
4
8
12
Time (weeks)
AC1 - Female
AC1 - Male
AC3 - Female
AC3 - Male
AAVrh32.33 NAb
AC1 - Female
AC1 - Male
AC3 - Female
AC3 - Male
AAV1/2/5/8/9 NAb
B
AC1 - Female
AC1 - Male
AC3 - Female
AC3 - Male
IFN-γ SFU / million PBMCs
150
100
50
0
0
4
8
Time (weeks)
No Stim.
AAVrh32.33 Pool A
AAVrh32.33 Pool B
AAVrh32.33 Pool C

### Figure S8

A

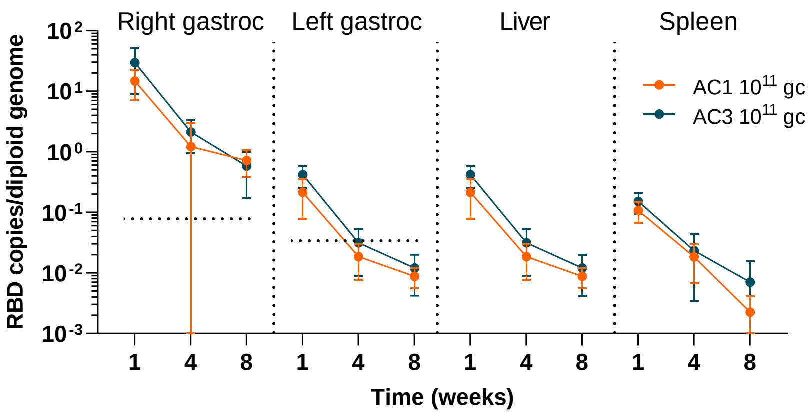


B

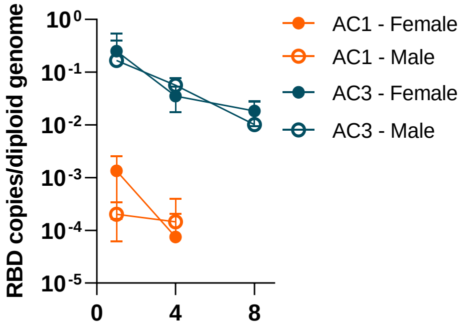


C

| | Time (weeks) | AC1 | | AC3 | |
|---|---|---|---|---|---|
| | | Mean | SD | Mean | SD |
| Right gastroc | 1 | 7.72E-04 | 8.92E-04 | 2.09E-01 | 2.18E-01 |
| | 4 | 1.10E-04 | 1.68E-04 | 4.58E-02 | 1.91E-02 |
| | 8 | ND | ND | 1.43E-02 | 1.22E-02 |
| Left gastroc | 1 | ND | ND | ND | ND |
| | 4 | ND | ND | ND | ND |
| | 8 | ND | ND | 1.53E-02 | 3.42E-02 |
| Liver | 1 | 5.09E-05 | 7.75E-05 | 1.47E-03 | 1.14E-03 |
| | 4 | ND | ND | 1.12E-04 | 2.51E-04 |
| | 8 | ND | ND | ND | ND |
| Spleen | 1 | ND | ND | 4.82E-05 | 1.08E-04 |
| | 4 | 3.20E-05 | 7.16E-05 | 2.07E-04 | 3.55E-04 |
| | 8 | 6.42E-05 | 1.44E-04 | 9.75E-05 | 2.18E-04 |

D

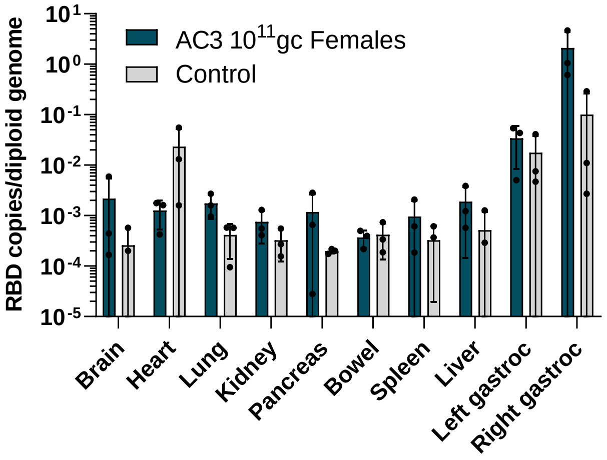

**Table S1. Related to Figures 5 and 6. Neutralizing AAV responses elicited by AAVCOVID in NHP.** Neutralizing antibody titers against the injected vector (AAVrh32.33) and cross-reactive neutralizing against other serotypes (AAV1, AAV2, AAV5, AAV8, AAV9) monthly after vaccination of rhesus macaques.

| **Animal** | **Time (weeks)** | **AAVrh32.33** | **AAV1** | **AAV2** | **AAV5** | **AAV8** | **AAV9** |
|---|---|---|---|---|---|---|---|
| **AC1 – female** | **0** | < 5 | < 5 | < 5 | < 5 | 5 | < 5 |
| | **4** | < 5 | < 5 | < 5 | < 5 | < 5 | < 5 |
| | **8** | < 5 | < 5 | < 5 | < 5 | < 5 | < 5 |
| | **12** | 20 | < 5 | < 5 | < 5 | < 5 | < 5 |
| | **14/16/20** | 80/80/160 | N/A | N/A | N/A | N/A | N/A |
| **AC1 - male** | **0** | < 5 | < 5 | < 5 | < 5 | < 5 | < 5 |
| | **4** | 10 | < 5 | < 5 | < 5 | < 5 | < 5 |
| | **8** | 160 | < 5 | < 5 | < 5 | < 5 | < 5 |
| | **12** | 320 | < 5 | < 5 | < 5 | < 5 | < 5 |
| | **14/16/20** | 320/320/320 | N/A | N/A | N/A | N/A | N/A |
| **AC3 - female** | **0** | < 5 | < 5 | < 5 | < 5 | < 5 | < 5 |
| | **4** | < 5 | < 5 | < 5 | < 5 | < 5 | < 5 |
| | **8** | 40 | < 5 | < 5 | < 5 | < 5 | N/A |
| | **12** | 160 | < 5 | < 5 | < 5 | < 5 | < 5 |
| | **14/16/20** | 160/160/160 | N/A | N/A | N/A | N/A | N/A |
| **AC3 - male** | **0** | < 5 | < 5 | < 5 | < 5 | < 5 | < 5 |
| | **4** | 5 | < 5 | < 5 | < 5 | < 5 | < 5 |
| | **8** | 40 | < 5 | < 5 | < 5 | < 5 | < 5 |

| | | | | | | | |
|---|---|---|---|---|---|---|---|
| | **12** | 80 | < 5 | < 5 | < 5 | < 5 | < 5 |
| | **14/16/20** | 320/320/160 | N/A | N/A | N/A | N/A | N/A |

**Table S2. Related to Figure 7. AAVCOVID stability assessment.** Titration (gc/mL) and percentage of titer relative to the initial titer of AC1 and AC3 aliquots stored at different temperatures for 1, 3, 7 or 28 days.

| | | Titer (gc/mL) | | Relative titer | |
|---|---|---|---|---|---|
| Temp | Time (days) | AC1 | AC3 | AC1 | AC3 |
| -80°C | NA | 2.65E+12 | 2.47E+12 | 100.00% | 100.00% |
| 4°C | 1 | 2.46E+12 | 2.23E+12 | 92.96% | 90.31% |
| | 3 | 2.30E+12 | 2.30E+12 | 86.91% | 93.01% |
| | 7 | 2.29E+12 | 2.40E+12 | 86.50% | 97.17% |
| | 28 | 2.49E+12 | 2.46E+12 | 93.96% | 99.31% |
| RT | 1 | 2.22E+12 | 2.33E+12 | 83.65% | 94.12% |
| | 3 | 2.33E+12 | 2.20E+12 | 88.05% | 89.03% |
| | 7 | 2.34E+12 | 2.39E+12 | 88.07% | 96.67% |
| | 28 | 2.37E+12 | 2.19E+12 | 89.35% | 88.75% |
